# The *Paracaedibacter*-like endosymbiont of *Bodo saltans* (Kinetoplastida) uses multiple putative toxin-antitoxin systems to maintain its host association

**DOI:** 10.1101/2020.07.24.217133

**Authors:** Samriti Midha, Daniel J. Rigden, Stefanos Siozios, Gregory D. D. Hurst, Andrew P. Jackson

## Abstract

Bacterial endosymbiosis has been instrumental in eukaryotic evolution, and includes both mutualistic, dependent and parasitic associations. Here we characterize an intracellular bacterium inhabiting the flagellated protist *Bodo saltans* (Kinetoplastida). We present a complete bacterial genome comprising a 1.39 Mb circular chromosome with 40.6% GC content. Fluorescent in situ hybridisation confirms that the endosymbiont is located adjacent to the nuclear membrane, and a detailed model of its intracellular niche is generated using serial block-face scanning electron microscopy. Phylogenomic analysis shows that the endosymbiont belongs to the Holosporales, most closely related to other α-proteobacterial endosymbionts of ciliates and amoebae. Comparative genomics indicates that it has a limited metabolism and is nutritionally host-dependent. However, the endosymbiont genome does encode diverse symbiont-specific secretory proteins, including a type VI secretion system and three separate toxin-antitoxin systems. We show that these systems are actively transcribed and hypothesize they represent a mechanism by which *B. saltans* becomes addicted to its endosymbiont. Consistent with this idea, attempts to cure *Bodo* of endosymbionts led to rapid and uniform cell death. This study adds kinetoplastid flagellates to ciliates and amoebae as hosts of *Paracaedibacter-like* bacteria, suggesting that these antagonistic endosymbioses became established very early in Eukaryotic evolution.

## Introduction

Eukaryotes commonly live in intimate associations with microbes (1). For microeukaryotes, intracellular microbes live as endosymbionts passing to progeny cells following fission (2), whereas endosymbionts of multicellular species live within host tissues and pass to progeny during reproduction, commonly inside eggs (3). Heritable host-microbe interactions have arisen multiple times and involve diverse eubacteria and life strategies.

The impact of endosymbioses on host and microbe vary widely. In some cases, the host captures a microbe for its own benefit. For instance, *Paramecium* captures *Chlorella* algae from the environment, utilizes them to facilitate a mixotrophic rather than heterotrophic lifestyle, but controls symbiont numbers such that *Chlorella* replication is higher when free-living than in the symbiotic state (4). In other cases, the microbe can be parasitic. This can occur when the symbiont is biparentally inherited, or uniparentally inherited, when it distorts reproduction towards the transmitting sex, for example with *Wolbachia* (5). Finally, and perhaps most commonly, symbiont and host mutually benefit from the interaction. For instance, symbionts such as *Buchnera* and *Euplotes* provide protective and nutritional benefits to their host that promote host survival and reproduction, and in doing so increase their fitness (6).

In many cases, host survival and reproduction depend on functions provided by the symbiont. For instance, loss of the anabolic capacity of *Buchnera* leaves their aphid host with insufficient essential amino acids that causes host sterility (6, 7). Host dependency may be due also to coadaptation of cellular and developmental processes (8). The defensive *Burkholderia* symbiont of the fungus *Rhizopus* is required for completion of the host’s sexual phase (9), a system mirroring the requirement of *Asobara tabida* wasps for one of their *Wolbachia* symbionts to complete oogenesis (10). In these cases, coadaptation over deep evolutionary time to the presence of the symbiont means that the host fails when the symbiont is removed, irrespective of any benefit the symbiont might confer.

There is another potential route to dependency that arises through selection on the symbiont to addict the host to its presence. Addiction mechanisms using toxin-antitoxin systems are a common retention mechanism for plasmids in bacteria (11) and are also observed in a variety of selfish genetic elements, such as the peel-zeel system of *C. elegans* (12). These systems typically function through delivery of a long-lived toxin molecule alongside an antitoxin with a shorter half-life. Should the element carrying the system not be inherited then the toxin becomes active and kills the non-bearer individual. Bacterial endosymbionts are currently not known to addict hosts through toxin-antitoxin systems, although they are known to exploit similar ‘rescue’ based systems in creating the phenotype of cytoplasmic incompatibility (12, 13).

In this study, we investigated the symbiosis between the flagellate *Bodo saltans* (Kinetoplastida) and its intracellular bacterial symbiont. *B. saltans* is a heterotrophic, phagocytic bacteriovore found in freshwater and marine habitats worldwide, and is among the closest free-living relatives of trypanosomatid parasites (14). Within the trypanosomatids, three parasitic lineages, *Novymonas, Phytomonas* and the Strigomonadinae, are known to contain a β-proteobacterial symbionts (15–17). These symbionts are mutualistic, cooperating in various metabolic pathways that provide essential amino acids and vitamins to the host (18–20). Although these endosymbionts are all mutualists, it is clear that parasitic kinetoplastids have acquired endosymbionts independently and relatively often (21). *B. saltans* is a free-living kinetoplastid, but microscopic examination has established that it too possesses endosymbiotic bacteria (22–24), whose precise identity is unknown.

We characterized the cytoplasmic bacterium inhabiting *B. saltans*, which we name *Candidatu*s Bodocaedibacter vickermanii, using a combination of genome sequencing, epifluorescence microscopy and serial block-face scanning electron microscopy (SBF-SEM). We then examined the endosymbiont genome sequence for putative function and identified a type VI secretion system and three chromosomal operons that are predicted to function as toxin-antitoxin systems. We therefore tested whether *Bodo* was dependent on its symbiont, potentially driven by these toxin-antitoxin systems, showing that antibiotic treatment results in rapid death of *B. saltans*, consistent with an addiction hypothesis.

## Material and Methods

### DNA sequencing and genome assembly

*Bodo saltans* cells were grown in 0.05% yeast extract in the presence of *Klebsiella pneumoniae* ATCC 13883 as prey. To exclude extracellular bacteria from the DNA extraction, cells were sorted by size using a BD FACS Aria cell sorter (Becton-Dickinson, USA). DNA was extracted from sorted cells using a Qiagen MagAttract HMW DNA Kit. DNA quality was assessed using a Qubit fluorometer (Thermo Fisher Scientific, USA) and Nanodrop instrument (Thermo Fisher Scientific, USA). A 20Kb insert library was prepared and sequenced on the PacBio Sequel platform (Pacific Biosciences). To assess *K. pneumoniae* contamination, reads were mapped using Bwa v0.7 to the published *B. saltans* genome sequence (CYKH01000000) and analysed with Samtools v0.1.18 (25). DNA sequence assembly was carried out using Canu v1.5 (26). To identify all distinct bacterial sequences present, rRNA reads were filtered using SortMeRNA v2.1 (27), which identified a single candidate endosymbiont. *De novo* assembly showed the presence of only one contig belonging to the same bacterium with terminal overlaps, indicating a complete circular chromosome. The endosymbiont genome sequence was circularized using the Amos package v3.1.0 (28) and constituent reads were mapped back to check for mis-assembly.

### Endosymbiont genome annotation

Genome annotation was carried out using Prokka v1.12 (29) complemented with BLASTp and InterProScan matches obtained using BLAST2GO v5.0 (30). Presence of prophage sequences were checked with PHAST (31). Genes containing a canonical signal peptide were predicted with SignalP 4.1 (32). Proteins with eukaryotic like domains were identified with EffectiveELD (33). Secretome P was used to predict genes encoding proteins secreted by non-classical pathway (34). Membrane transporters were predicted using TransportTP (35) and transporters were classified using the BLAST tool of the Transport Classification DataBase (36). HHsearch (37) was used to search for distant relationships between putative toxin-antitoxin system components and known structures at the Protein Data Bank (PDB;(38)) or protein families in the Pfam database (39). Three iterations of jackhmmer (40) searching against the UniClust database (41) were used to generate the sequence profiles for comparison with database entries. 16S rRNA sequences were predicted using RNAmmer v1.2 and EzBioCloud was used for taxonomical identification of the bacterial lineage (42).

### Phylogenetic analysis and comparative genomics

OrthoMCL v2.0.9 was used to cluster orthologous genes shared by the *Bodo* endosymbiont with 12 related alpha-proteobacterial genome sequences (43). Protein sequences encoded by 187 conserved genes from the 14 bacterial genomes were aligned using ClustalW (44). Prottest v3.0 was used to check for the optimal amino acid substitution model for phylogeny estimation (45). An LG+I+G model was applied in RaxML to estimate a maximum likelihood tree with 1000 non-parametric bootstrap replicates (46). BLAST-based Average Amino Acid Identity (AAI) values amongst the 13 genomes were calculated using AAI calculator (47). MAPLE 2.3.0 was used to identify and compare the completeness of various metabolic pathways in these endosymbiont genomes (48). Genomaple 2.3.2 was used to compare the metabolic pathways of endosymbiont and *B. saltans* (48).

### Fluorescent in situ hybridization (FISH)

The endosymbiont 16S rRNA sequence was integrated into the SILVA database using the web-based SINA aligner (49, 50). This alignment was further merged with SILVA Ref NR 99 (51) using the ARB package and the probe design tool was used to select FISH probes specific for the endosymbiont genome. (52). The probes were tested for mismatch analysis using mathFISH, probeCheck and BLAST (53, 54). *B. saltans* cells were fixed using 4% paraformaldehyde and spotted on 0.1 % gelatin coated slides. Slides were dehydrated with 50%, 80% and 100% ethanol consecutively before hybridization with the probe (50 ng/μl) at 46°C. The probe was labelled with cyanine dye (([CY3]CGAAGTGAAATCTACGTCTCCGT)) and hybridized with 15% formamide in three independent replicates. Counterstaining of the cells was achieved using VECTASHIELD Antifade Mounting Medium with DAPI.

### Electron microscopy

Cells were fixed in 2.5% glutaraldehyde (Wt/Vol) in 0.1M phosphate buffer (pH7.4) in Pelco Biowave (Ted Pella Inc.) and washed twice in 0.1M PB before embedding in 3% agarose. Agarose embedded cell pellets were post-fixed and stained as described previously (55), except for use of 0.1% thiocarbohydrazide as a mordant. For TEM, after UA staining samples were embedded with TAAB medium Premix resin in silicone moulds and Beem capsules. Ultrathin serial section (70-75nm) were cut on an UC6 ultra-microtome (Leica, Vienna) collected on formvar coated copper grids, before viewing at 120KV in a FEI Tecnai G2 Spirit. Images were taken using a MegaView III camera and multiple Image Alignment (MIA) was used to create a high-resolution overview of areas of interest.

For SBF-SEM, cell pellets were embedded with TAAB hard premix resin in plastic dishes. Excess resin was removed before the block was mounted onto a cryo pin, cell side up, using silver conductive epoxy. Targeted trimming created a block face of cells 500μm x 500μm. Samples were painted and dissected as previously described (55). To mitigate charge build up and maximize image quality, imaging conditions were as follows: low vacuum mode with a chamber pressure of 70 Pa. Low accelerating voltage (1.7kV), dwell time per pixel (14μs), magnification (5040×) pixel size 5.4 nm in x and y, frame width 6144 × 6144, section thickness 100 nm over 104 sections. Amira 6.5.0 was used to analyse the SBF-SEM images and generate a 3D model for the cell.

### Attempt to cure the symbiont with Rifampicin treatment

Besides *B. saltans*, two parasitic and asymbiotic kinetoplastids, *Trypanosoma theileri* and *Leptomonas costaricensis*, were treated with rifampicin (20 μg/ml), alongside control populations. To avoid impacts of rifampicin on *Bodo* mediated through changes in bacterial prey abundance, a rif-resistant prey strain of *Klebsiella* was developed for these experiments. The number of kinetoplastid cells was counted at 0 hours and after 24 hours of treatment in control and treated flasks. The experiment was repeated three times and statistical significance of changes in cell number between species were analysed using ratio paired T-test.

### Bodo saltans RNA-sequencing

*B. saltans* cells were treated with gentamicin (50 μg/ml) before RNA extraction. Total RNA was extracted from cells and rRNA was depleted using RiboZero rRNA depletion kit (Epidemiology). A strand-specific library was constructed with the NEBNext Ultra Directional RNA library preparation kit (NEB). Paired end 2×2013;150 bp sequencing was carried out on the Illumina platform. Raw reads were mapped on to the endosymbiont genome sequence using bwa v0.7.12 and low quality read alignment were removed using samclip (https://github.com/tseemann/samclip) and default parameters. Sort read alignments were visualised with the IGV 2.8.6 tool, while strand-specific read quantitation was performed with the SeqMonk program (https://www.bioinformatics.babraham.ac.uk/proiects/seqmonk/). To account for the presence of low DNA contamination in our library, a Difference Quantitation correction was applied. Read counts for each gene were calculated as the read counts originated from the opposite strand (template strand) minus the read counts from the same strand (coding strand) and normalized to the length of each gene (RPK).

### Data availability

The *Candidatus* Bodocaedibacter vickermanii genome assembly has been submitted to NCBI under accession number CP054719. Raw PacBio genomic reads and Illumna RNA-Seq reads were submitted to Sequence Read Archive (SRA) database under the accession number SRR11932788 and SRR11935165 respectively.

## Results

### A *Bodo saltans* endosymbiont of the Paracaedibacteraceae

A single PacBio Sequel sequencing run generated 697,281 reads; of these, ~78% mapped to the *B. saltans* genome. Multiple proteobacterial genomes were discovered among the assembled contigs (Supplementary figure 1). Analysis of prokaryotic rRNA reads identified four bacterial taxa; *Klebsiella*, *Cupriavidus*, *Delftia* (each 100% identical with environmental sequences and assumed to derive from prey bacteria in the cell culture), and an unknown bacterium with greatest sequence identity (98.59%) to uncultured bacterial sequences belonging to Family Paracaedibacteraceae, followed by *Paracaedibacter* (86.89%). Given that this fourth taxon has such low sequence identity with any defined genus, we propose that it represents the endosymbiont and hereafter refer to it as *Candidatus* Bodocaedibacter vickermanii (Cbv) gen. nov. sp. nov.. The organism is named after Keith Vickerman (1933-2016), author of seminal microscopic studies of diverse kinetoplastids, including *B. saltans*.

### General features of the *Candidatus* Bodocaedibacter vickermanii genome

The genome containing the *Candidatus* Bodocaedibacter vickermanii rRNA sequence was assembled as a complete chromosome, 1.39 Mb in size and with an average coverage of 212x. The GC skew graph (Fig. 1A) shows the pattern typical of a complete bacterial chromosome, transitional points referring to the origin and terminus of replication. The genome is 40.7% GC in content and encodes 1214 putative CDS, 40 tRNA and 2 rRNA-encoding operons (both rRNA operons have 16S and 23S rRNA genes in close vicinity, separated by two tRNAs). The genome also possesses two incomplete prophage elements (CPBP_00240-CPBP_00250; CPBP_00316-CPBP_00324).

**Figure 1:**
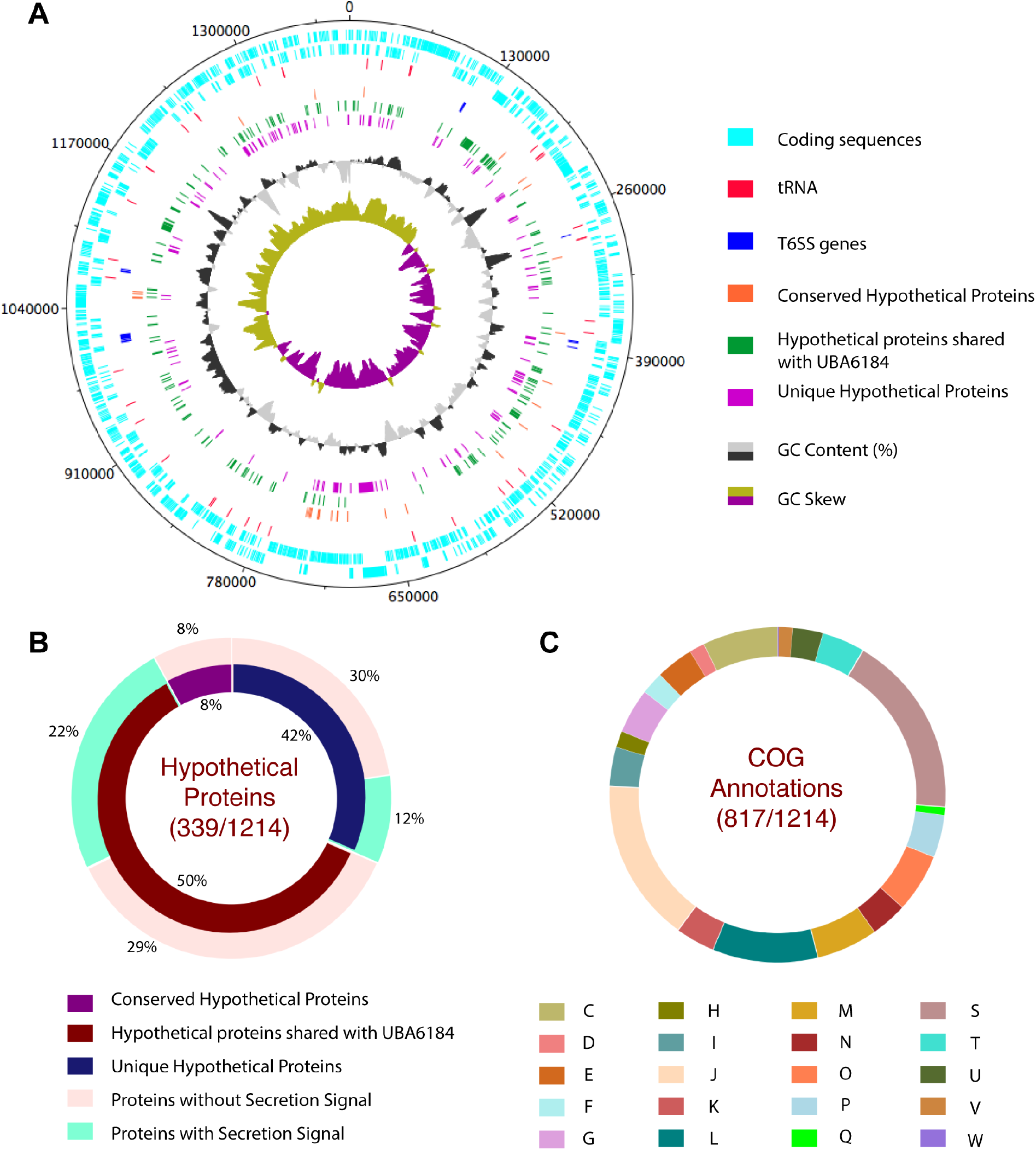
Annotation features of Bodo endosymbiont. **A.** Graphical representation of *Candidatus* Bodocaedibacter vickermanii (Cbv) genome, generated using DNAPlotter v10.2 (86). Moving inwards, the tracks represent forward and reverse CDS, tRNAs, type VI secretion system (T6SS) encoding genes, hypothetical proteins (conserved in the *Candidatus* Paracaedibacteraceae bacterium UBA6184 genome and unique to Cbv), GC (%) plot and GC skew [(G-C)/(G+C)] plot. **B.** Percentage of conserved and unique hypothetical proteins encoded in genome and presence of secretion signal is shown in pie chart. **C.** Cluster of Orthologous Genes (COG) annotation categorization is shown in pie chart. Categories: Energy production and conversion (*C*), Cell cycle control, cell division, chromosome partitioning (*D*), Amino acid transport and metabolism (*E*), Nucleotide transport and metabolism (*F*), Carbohydrate transport and metabolism (*G*), Coenzyme transport and metabolism (*H*), Lipid transport and metabolism (*I*), Translation, ribosomal structure and biogenesis (*J*), Transcription (*K*), Replication, recombination and repair (*L*), Cell wall/membrane/envelope biogenesis (*M*), Cell Motility (*N*), Post-translational modification, protein turnover, chaperone functions (*O*), Inorganic ion transport and metabolism (*P*), Secondary metabolites biosynthesis, transport and catabolism *(Q)*, Function Unknown (*S*), Signal transduction mechanisms *(T)*, Intracellular trafficking, secretion, and vesicular transport (*U*), Defence mechanisms (*V*), Extracellular structures (*W*).

Among sequenced genomes (Table 1), the *Ca*. B. vickermanii genome sequence is most closely related to a metagenomics sequence assembly from Canadian waste water (UBA6184) (56), and thereafter to various endosymbiont genomes of the Holosporales (86-87% 16S rRNA sequence identity). 27% of gene sequences in the *Bodo* symbiont genome encode uncharacterized proteins found mostly in *Ca*. B. vickermanii and the UBA6184 metagenome (Fig. 1B). A majority (~67%) of putative coding sequences were placed into one of 20 functional categories of Clusters of Orthologous Groups of proteins (COG), shown in Fig. 1C, while 18% of annotated proteins were categorized as having ‘unknown function’.

**Table 1:** List of genomes used in the study: Annotation features, isolation source and accession numbers

| S.No. | Organism | GC content (%) | Size (Mb) | CDS | tRNA | rRNA | Family | Order | Phylum | Host/Isolation source | Accession number |
| --- | --- | --- | --- | --- | --- | --- | --- | --- | --- | --- | --- |
| 1 | *Candidatus* Caedibacter acanthamoebae 30173 | 38.2 | 1.72 | 1359 | 42 | 3 | Caedimonadaceae | Holosporales | Proteobacteria | *Acanthamoeba polyphaga* HN-3 | CP008936 |
| 2 | *Caedibacter varicaedens* | 42.1 | 1.69 | 1726 | 41 | 3 | Caedimonadaceae | Holosporales | Proteobacteria | *Paramecium bicaurelia* | BBVC00000000 |
| 3 | *Candidatus* Paracaedibacter symbiosus PRA9 | 41.5 | 2.66 | 2199 | 43 | 6 | Paracaedibacteraceae | Holosporales | Proteobacteria | *Acanthamoeba* sp. UWET39 | JQAK00000000 |
| 4 | *Candidatus* Odysella thessalonicensis L13 | 42 | 2.85 | 2398 | 40 | 3 | Paracaedibacteraceae | Holosporales | Proteobacteria | *Acanthamoeba* | AEWF00000000 |
| 5 | *Candidatus* Paracaedibacter acanthamoebae PRA3 | 41.0 | 2.47 | 2154 | 43 | 6 | Paracaedibacteraceae | Holosporales | Proteobacteria | *Acanthamoeba* | CP008941 |
| 6 | *Candidatus* Bodocaedibacter vickermanii | 40.7 | 1.39 | 1214 | 40 | 6 | Paracaedibacteraceae | Holosporales | Proteobacteria | *Bodo saltans* | CP054719 (this study) |
| 7 | *Candidatus* Paracaedibacteraceae bacterium UBA6184 | 40.1 | 1.47 | 1444 | 30 | - | Paracaedibacteraceae | Holosporales | Proteobacteria | waste water | DITM00000000 |
| 8 | *Holospora curviuscula* 02AZ16 | 38.1 | 1.72 | 1534 | 42 | 3 | Holosporaceae | Holosporales | Proteobacteria | *Paramecium bursaria* | PHHC00000000 |
| 9 | *Holospora obtusa* F1 | 35.2 | 1.33 | 1249 | 36 | 3 | Holosporaceae | Holosporales | Proteobacteria | *Paramecium caudatum* | AWTR000000000 |
| 10 | *Holospora undulata* HU1 | 36.1 | 1.40 | 1460 | 44 | 3 | Holosporaceae | Holosporales | Proteobacteria | *Paramecium caudatum* | ARPM00000000 |
| 11 | *Holospora elegans* E1 | 36.0 | 1.27 | 1454 | 37 | 3 | Holosporaceae | Holosporales | Proteobacteria | *Paramecium caudatum* | BAUP00000000 |
| 12 | Endosymbiont of<br>*Acanthamoeba* UWC8 | 34.8 | 1.62 | 1494 | 36 | 3 | Candidatus<br>Midichloriaceae | Rickettsiales | Proteobacteria | *Acanthamoeba* | CP004403 |
| 13 | *Candidatus* Jidaibacter<br>acanthamoeba UWC36 | 33.7 | 2.37 | 2267 | 36 | 3 | Candidatus<br>Midichloriaceae | Rickettsiales | Proteobacteria | *Acanthamoeba*<br>*sp. UWC36* | JSWE00000000 |
| 14 | *Magnetococcus marinus*<br>MC-1 | 54.2 | 4.72 | 3766 | 45 | 9 | Magnetococcaceae | Magnetococcal<br>es | Proteobacteria | marine water | CP000471 |

### Microscopy indicates this microbe represents an intracellular symbiont

To confirm that the *Ca*. B. vickermanii sequence is correctly attributed to an intracellular endosymbiont, we used Fluorescent *in situ* Hybridization (FISH) and *Ca*. B. vickermanii 16S rRNA sequence probes. Cy-3 labelled probes bound to bacterial DNA on fixed *B. saltans* cultures, which were visualized using DIC microscopy (Fig. 2A). Stained *B. saltans* nuclei and kinetoplast are seen in blue channel (Fig. 2B) and bacterial spots are seen in red channel (Fig. 2C). A merged image from (Fig. 2D) shows that the bacteria are found adjacent to the *B. saltans* nucleus and are not observed extracellularly.

**Figure 2:**
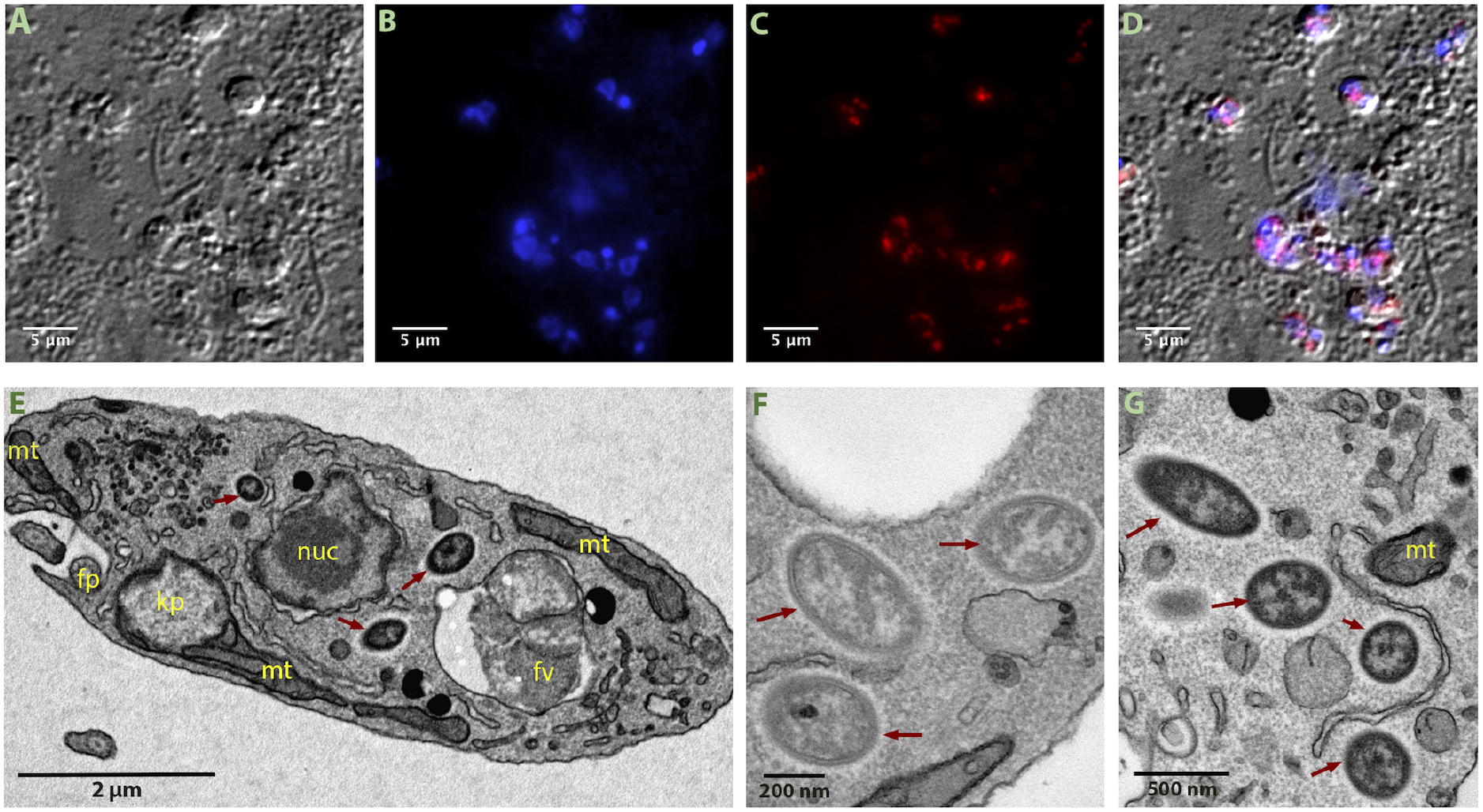
Visualization of intracellular bacteria in *Bodo saltans*. Images from Fluorescent *in situ* hybridization (FISH) experiment (A-D) and transmission electron microscopy (TEM) experiment (E-G). Imaging of *B. saltans* cell cultures with differential interference contrast (A), staining with DAPI to visualize nucleus and kinteoplast (B), FISH staining with endosymbiont specific probe conjugated to Cy3 (C) and images from three channels overlaid (D). Ultrastructure of the *B. saltans* cell (E) displaying nucleus (nuc), kinetoplast (kp), mitochondria (mt), food vacuole (fv), flagellar pocket (fp) and three intracellular bacteria marked with dark red arrows, endosymbionts showing the presence of an electron lucid halo around cell membrane (F-G). Images were acquired on a Zeiss Axio Observer Z1 (Carl Zeiss AG, Jena, Germany) equipped with 100x 1.4NA objective, 2.5x optovar. Images captured were analyzed using ImageJ v2.0 (87).

Electron microscopy was used to further define the intracellular niche. In the TEM images (Fig. 2E), a large nucleus surrounded by double membrane occupies the cell centre. The kinetoplast is positioned consistently adjacent to basal bodies. Multiple mitochondrial sections are seen along the cellular periphery. Food vacuoles, enclosing engulfed bacteria, are usually evident towards the posterior end. In addition to these typically kinetoplastid features, we identified rod-shaped bacteria-like structures that are 0.9-1.2 μm in length and 0.3-0.4 μm in diameter and often surrounded by an electron lucid halo (Fig. 2F-2G). These are consistent with endosymbiont cells previously observed in *B. saltans* (23, 24).

3-D models generated from SBF-SEM imaging further resolved the disposition of these rod-shaped structures. Fig. 3 shows the bean-shaped *B. saltans* cell with characteristic kinetoplastid features. In this 3-D rendering, rod-shaped bacterial cells are free in the cytoplasm, close to the nucleus. The number of bacterial cells varied from 3 to 10 (N=40). Where a higher bacterial count was observed, a membrane enveloping those cells adjacent to the *Bodo* nucleus was often seen, similar to that previously observed (24). This could be the nuclear membrane enveloping the endosymbionts prior to cell division and distribution into daughter cells.

**Figure 3:**
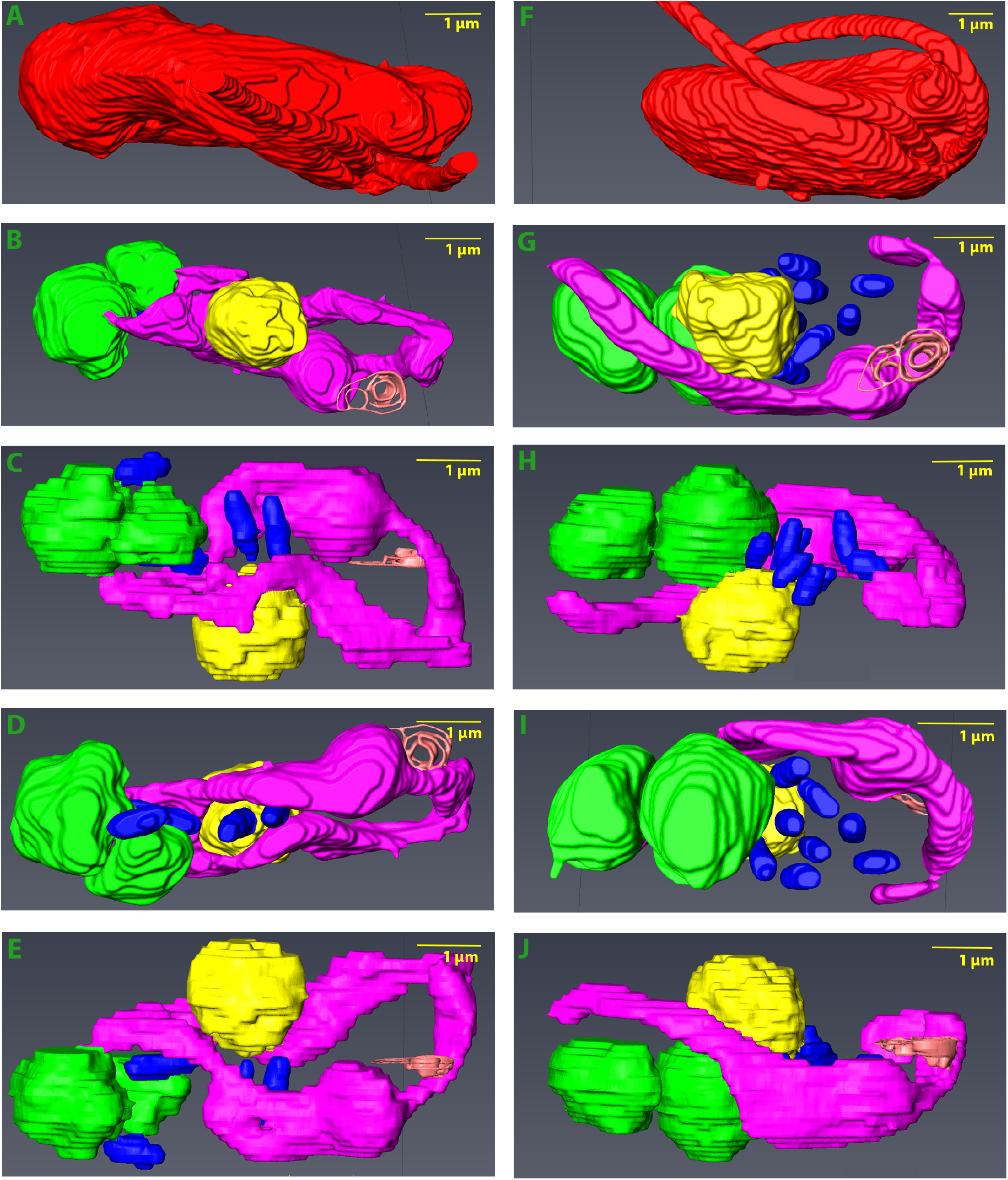
3D model of *Bodo saltans* cell generated using Serial Block Face-Scanning Electron Microscopy. Animations of *B. saltans* cellular ultrastructure in longitudinal section. Two cells are shown with differences in endosymbiont number and distribution: cell 1 (A-E) and cell 2 (F-J). For each cell, one image with cell envelope (A, F) and four longitudinal sections with consecutive 90-degree rotation around radial axis are shown (B-E and G-J). Cell envelope (in red) shows the bean shaped structure of *B. saltans* and two flagella emerging from flagellar pocket (in orange). Nucleus (in yellow) is situated in the centre of the cell and cytobionts (in blue) are distributed in close vicinity to the nucleus. A large, swirled mitochondrion (in magenta) is placed around the periphery of the cell, with a kinetoplast capsule just under the flagellar pocket. Multiple food vacuoles (in green) are present at the posterior end of the cell.

### *Ca*. Bodocaedibacter is a novel genus in a clade of alpha-proteobacterial endosymbionts

A maximum likelihood phylogenetic tree was estimated from an alignment of 187 single copy core genes from 13 genomes belonging to alpha-proteobacterial endosymbionts of protists of families Paracaedibacteraceae, Caedimonadaceae, Midichloriaceae and Holosporaceae (Table 1). The genome of *Magnetococcus marinus*, belonging to basal lineage of alpha-proteobacteria, was used as an outgroup. The tree places the *Bodo* endosymbiont among other Holosporales and Rickettsiales endosymbionts of amoebae and ciliates, forming a clade with *Holospora, Caedibacter* and *Paracaedibacter-like* organisms (Fig. 4). As with the 16S rRNA homology analysis, the *Ca*. B. vickermanii genome grouped most closely with a wastewater metagenome (UBA6184). Branch lengths are long, consistent with these higher-level taxonomic comparisons. We examined the average amino acid identity (AAI) values between the *Bodo* endosymbiont and the 13 related genomes (Fig. 4) and, with the exception of UBA6184, AAI values for all related genomes are lower than the conventional genus boundary of 55% (57), indicating that *Ca*. B. vickermanii and UBA6184 represent a novel genus.

**Figure 4:**
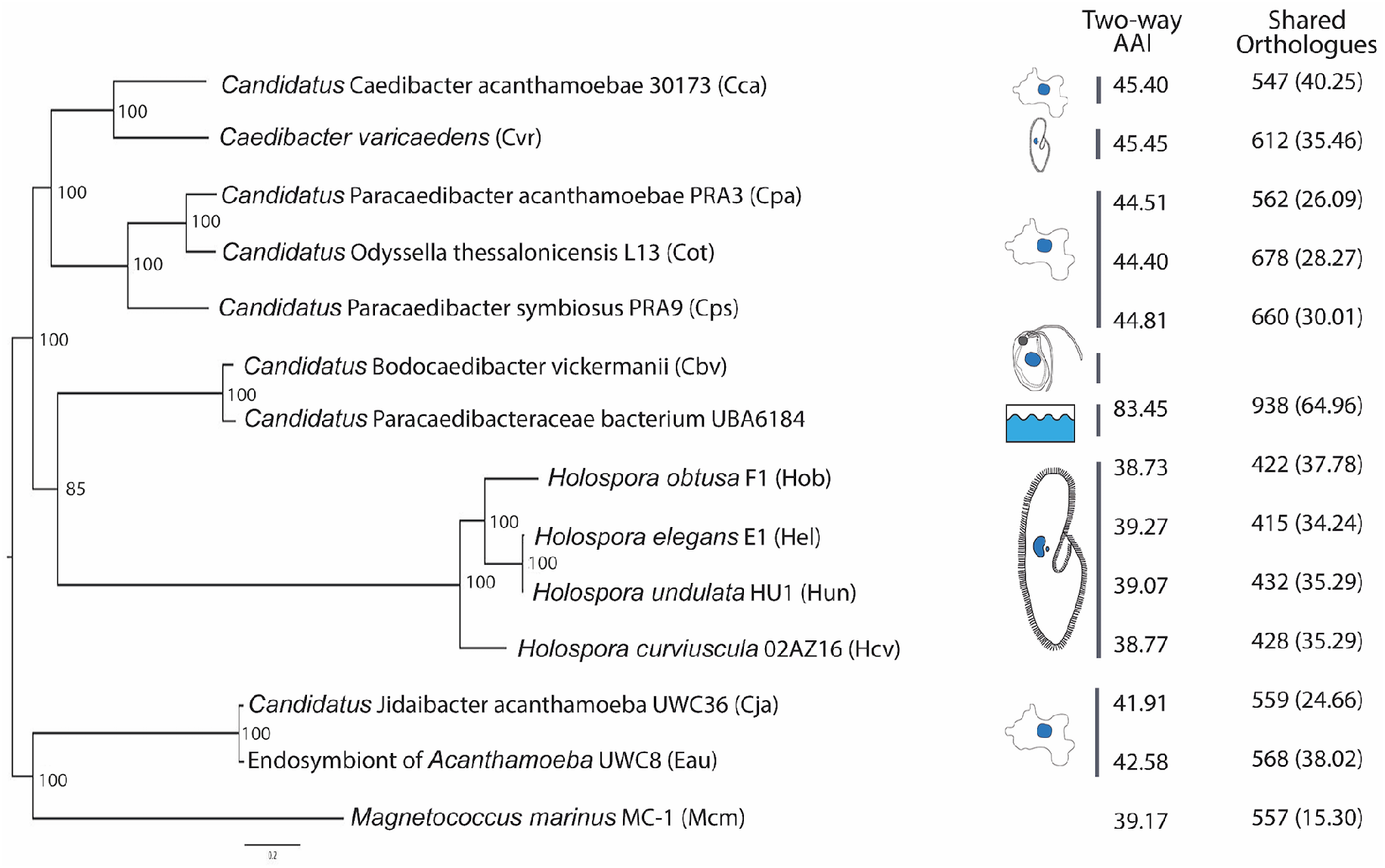
Phylogenetic relationships of *Candidatus* Bodocaedibacter vickermanii and other alpha-proteobacterial endosymbionts of protists. A maximum likelihood tree estimated from a concatenated alignment of 187 protein sequences. The tree is rooted with an outgroup (Mcm). Bootstrap values shown on the nodes are calculated from 1000 non-parametric replicates. The scale bar (0.3) is the number of amino acid substitutions per site. Cartoons shown next to tree represent the isolation source (see Table 1 for details). Two columns on the right side depict the average amino acid identity (AAI) and number of orthologous gene clusters shared between *Ca*. B. vickermanii and each other genome sequence, with the corresponding value as a percentage of all genes shown in brackets.

Analysis of gene content using OrthoMCL (38) shows that these 13 endosymbiotic genomes share a core repertoire of 242 genes, and typically <50% of their total gene repertoire (Fig. 4). Mapping these gene sets to KEGG pathways indicates that many are involved in central metabolism and information processing (Supplementary figure 2).

### The *Ca*. B. vickermanii genome has limited metabolic capacity

The KEGG classifications of coding sequences in the *Ca*. B. vickermanii genome and 12 related endosymbiont sequences show that the *Bodo* endosymbiont has a small genome and relatively limited metabolic capacity (Table 1). We calculated Module Completion Ratios (MCR) for purine metabolism, pyrimidine metabolism, amino acid metabolism, polyamine biosynthesis, cofactor and vitamin biosynthesis pathways using MAPLE (48) (Supplementary Fig. 3). While often present in related genomes, several of these pathways are either completely or partially absent in *Ca*. B. vickermanii. For example, it lacks biotin, pantothenate and coenzyme A biosynthesis pathways entirely, and encodes only five out of nine genes in the ubiquinone biosynthesis pathway. Conversely, *Ca*. B. vickermanii possesses diverse membrane transporter genes for metabolite acquisition. The TransportTP (35) server identified 81 genes encoding membrane transporters in *Ca*. B. vickermanii, more than *Holospora* genomes (56–67) but rather fewer than other *Caedibacter/Paracaedibacter* genomes (102-148).

A full account of all transporter genes, classified using the Transporter Classification Database (36), is shown in Supplementary Table 1. These include genes associated with amino acid transport, metabolite transport (e.g. Drug/Metabolite Transporter (DMT) family (n=2)), the Major Facilitator Superfamily (MFS) (n=22), and the ATP-binding Cassette (ABC) superfamily (n=27). There are also genes encoding for exporter proteins like the Resistance-Nodulation-Cell Division (RND) Superfamily (n=6) and the Multidrug/Oligosaccharidyl-lipid/Polysaccharide (MOP) Flippase Superfamily (n=4).

We also examined the possibility that metabolic pathways are conducted cooperatively, with component genes being drawn from both host and endosymbiont genomes, which might indicate a mutualism. Only two pathways, lysine and threonine biosynthesis, appeared as potential candidates for co-operation, as shown in Supplementary figure 4. These two instances aside, most of the pathways associated with essential amino acids, vitamins and cofactor biosynthesis were lacking in both organisms, quite unlike the mutualistic *Novymonas* and Strigomonadinae endosymbiosis (18–20), indicating that *B. saltans* remains firmly heterotrophic (Supplementary figure 5). Taken together, this does not suggest that *Ca*. B. vickermanii provides obvious metabolic benefit to *B. saltans*; indeed, the endosymbiont appears to be nutritionally dependent on its host.

### The *Ca*. B. vickermanii genome encodes a Sec pathway and Type VI Secretion System (T6SS)

We examined the endosymbiont genome for various secretion systems that could facilitate communication with the host. This gram-negative bacterium contains the cellular components for a specific Type VI Secretion System (T6SS) and the general secretion (Sec) pathway. The *Ca*. B. vickermanii genome encodes most of the essential T6SS components, namely genes associated with membrane complex (tssL and tssM), baseplate complex (tssA, tssE, tssF, tssG, tssK and tssI/VgrG) and tail complex (tssB and tssC). Analysis with InterproScan also identified a tssJ-like gene (CPBP_00933) encoding a T6SS-associated lipoprotein and a gene (CPBP_00987) encoding a putative Hcp-like superfamily protein. RNA-seq analysis confirmed that these genes are actively transcribed, albeit at low levels (Supplementary table 2).

In Paracaedibacteraceae and Caedimonadaceae genomes T6SS core genes are organised at four or more different genomic locations in a conserved arrangement: [tssG-tssF], [tssB-tssC-hcp], [tssK-tssL-tssM-tssA], and [ywqK-vgrG]. The *Ca*. B. vickermanii genome complies with this convention (Supplementary Fig. 6), indicating that the system could be functional in *Ca*. B. vickermanii, as in related endosymbionts (58–61).

The web-based server Bastion6 was used to predict for proteins potentially secreted by the T6SS (62). This machine learning-based algorithm identified 117 proteins as putative type VI secretion effectors (T6SE) (Supplementary table 3), which included various enzymes, flagellar proteins, membrane proteins and numerous uncharacterized proteins.

The Sec pathway is a major pathway for protein translocation across the cell membrane (63). The *Ca*. B. vickermanii genome includes genes for a putative motor protein SecA, translocase complex SecYEG, and the auxillary components SecDF-YajC and YidC. It also encodes the proteins required for co-translational targeting (signal recognition particle proteins and receptor FtsY), as well as post-translational targeting (protein targeting component SecB). To discover proteins that may be targeted by the Sec pathway for integration or secretion outside the cell, we identified 158 coding sequences containing a signal sequence using SignalP 4.1 (32) (Supplementary Table 4). Of these, 105 proteins (66.4%) are uncharacterized and only 21 contain predicted transmembrane helices, indicating that most are probably secreted.

The genes predicted to encode for proteins with a secretion signal include various enzymes (xylanase, serine endoprotease, proteases, phospholipase, methyltransferase and acetyltransferase), flagellar proteins and membrane transporters. They also include two genes encoding proteins containing tetratricopeptide-repeats, and one gene for ankyrin-repeat containing protein, which are both typical eukaryotic protein domains involved in protein-protein interactions. A majority of these genes (130/158) have homologs only in the UBA6184 genome. The 105 genes encoding uncharacterized proteins with secretory signals are also limited to these two bacterial genomes, indicating this lineage has evolved a considerable repertoire of specific secreted proteins.

In case non-classical pathways are used to secrete *Ca*. B. vickermanii proteins, we analysed predicted protein sequences with SecretomeP (34). There are 204 proteins that returned a score above the threshold, 59 of which also returned a significant result with SignalP (Supplementary Table 4). Also among these cases were genes encoding proteins with homology to known cellular toxins, which led to the discovery of three putative novel toxin-antitoxin islands in the *Bodo* endosymbiont.

### Multiple polymorphic toxin systems are a possible mechanism for addiction

Bacterial Polymorphic Toxin Systems (PTS) such as the Rhs (64) and CDI (65) systems comprise genes encoding toxins and their cognate anti-toxins in characteristic operons (66). Typically, a toxin gene encodes a large multi-domain protein contains an N-terminal secretion signal followed by a toxin domain. A toxin gene is immediately followed by a gene encoding an anti-toxin, (or ‘immunity protein’), capable of neutralising the toxin. This pair is often followed by additional ‘orphan modules’, each encoding an alternative toxin with its cognate antitoxin. Genes in these orphan modules may often contain a repeat region homologous to part of the toxin gene at the beginning of the operon. This repeat enables recombination between the full-length toxin gene and an orphan module to generate an alternative toxin protein.

We identified three genomic regions coding for putative PTS (Fig 5; Table 2). The putative toxin proteins identified here have no structural homology with known proteins, nor is there any homology between the three putative PTS. Nevertheless, the three regions show typical PTS characteristics. These include the presence of regions of homology between putative toxin proteins in the N-terminal part preceding the toxin domains (Fig 5) and the organisation of putative toxin and antitoxin proteins across the loci (Table 2), which leads us to conclude that they may represent novel PTS. In many cases, the relationships used to infer toxin or antitoxin function are characterised by low degrees of sequence identity, but the HHsearch (37) probability scores are all >80% (and often much higher), which supports genuine homology.

**Figure 5:**
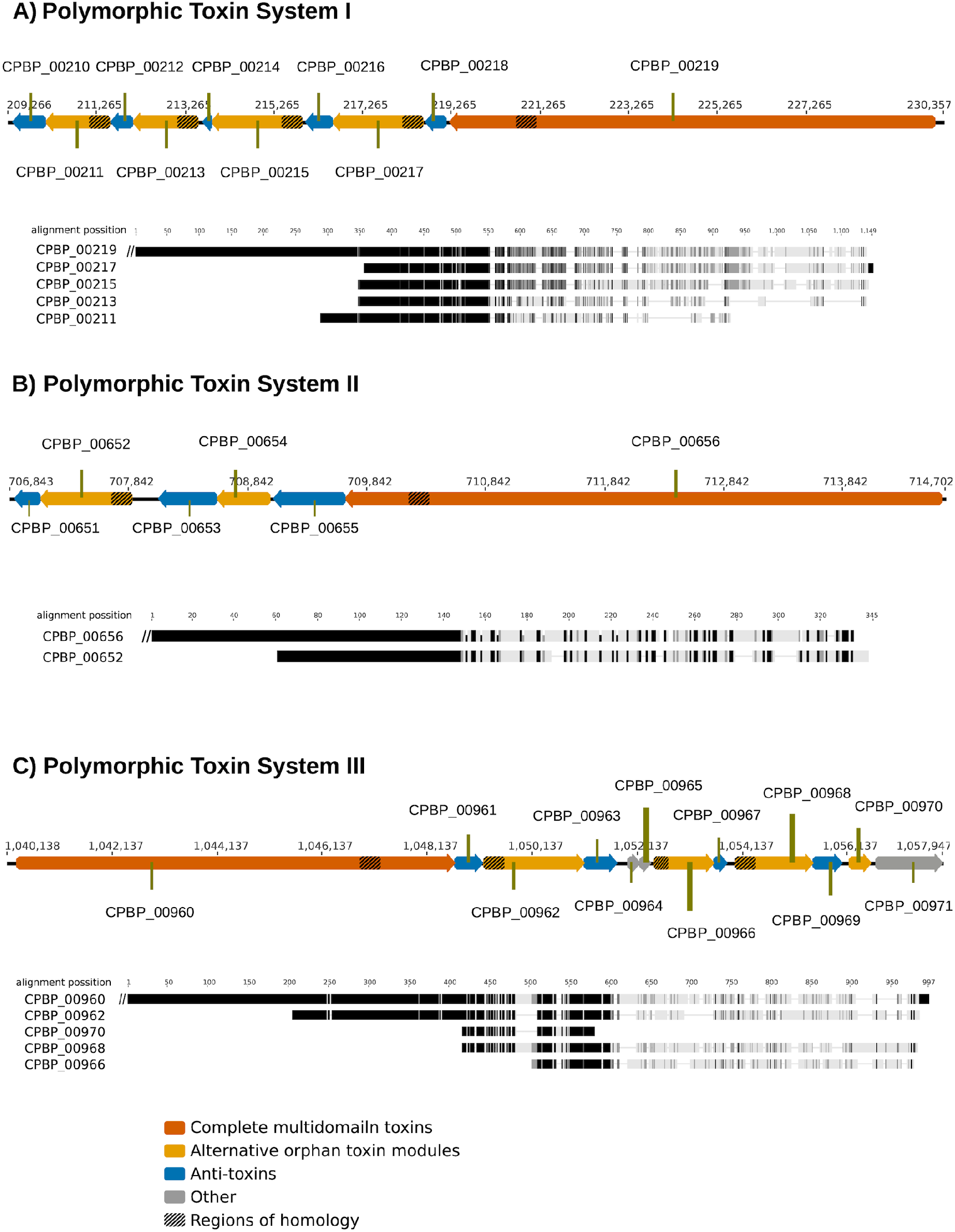
Polymorphic toxin systems in *Ca*. B. vickermanii. Schematic diagrams of the organization of the three predicted polymorphic toxin systems in *Ca*. B. vickermanii. In each case, the putative, alternative orphan toxin modules are shown downstream of the main multi-domain toxin. Beneath each is a protein multiple sequence alignment of the main toxin C-terminus and the alternative orphan sequences, generated using MAFFT program under default parameters as implemented in Geneious software v6.7. Gray scale gradient represents identical (black), similar (dark gray) and dissimilar (light gray) residues in the alignment based on the Blosum62 score matrix. Note that the specification of orphan modules is least certain for PTS III. Although CPBP_00970 resembles the N-terminus of CPBP_00960, its coding sequence is abbreviated. This, and the metalloprotease (PF03410) match of CPBP_00971 that would, by expected PTS gene positioning encode an antitoxin, argue against this pair of genes representing an orphan module. Another pair of genes that should, by the PTS gene positioning scheme, be an orphan module are CPBP_00964 and CPBP_00965. However, these two sequences are short (75 and 67 residues respectively), homologous (around 34% identical) and CPBP_00964 lacks any homology to the N-terminus of CPBP_00960, arguing against these genes constituting an orphan module.

**Table 2:** Annotation of polymorphic toxin systems by distant homology detection with HHsearch

| Gene | Module position | Description based on position and/or matching families | Matching Pfam or PDB entry suggestive of toxin or antitoxin function | Matching probability (%), sequence identity of HHsearch alignment |
| --- | --- | --- | --- | --- |
| *Polymorphic Toxin System I* |  |  |  |  |
| CPBP_00219 |  | Complete multidomain toxin | AHH nuclease domain (PF14412) | 93, 23 |
| CPBP_00218 |  | Antitoxin | Gmx_para_CXXCG (PF09535); DUF1629 (PF07791); Immunity protein 43 (PF15570) | 100, 14;<br>99, 15;<br>96, 16 |
| CPBP_00217 | Orphan module 1 | Alternate toxin domain | Tox-HNH-HHH nuclease domain (PF15637) | 100, 34 |
| CPBP_00216 |  | Antitoxin | SUKH superfamily 5 (PF14567) | 88, 15 |
| CPBP_00215 | Orphan module 2 | Alternate toxin domain | Cloacin; Colicin-like bacteriocin tRNase domain (PF03515) | 99, 6 |
| CPBP_00214 |  | Antitoxin | Colicin/pyocin immunity protein (PF01320) | 100,16 |
| CPBP_00213 | Probable orphan module 3 | Alternate toxin domain | *No strong match to any family or structure* |  |
| CPBP_00212 |  | Antitoxin | *No strong match to any family or structure* |  |
| CPBP_00211 | Orphan module 4 | Alternate toxin domain | DUF1837 (PF08878) | 81, 16 |
| CPBP_00210 |  | Antitoxin | Immunity protein 49 (PF15575) | 100,12 |

| ***Polymorphic Toxin System II*** |  |  |  |  |
| --- | --- | --- | --- | --- |
| CPBP_00656 |  | Complete multidomain toxin | Tox-HNH-HHH nuclease domain (PF15637) | 99, 40 |
| CPBP_00655 |  | Antitoxin | SUKH superfamily 5 (PF14567) | 91, 15 |
| CPBP_00654 | Orphan module 1 | Alternate toxin domain | DNase/tRNase domain of colicin-like bacteriocin (PF12639) | 100, 28 |
| CPBP_00653 |  | Antitoxin | SUKH superfamily 5 (PF14567) | 100, 12 |
| CPBP_00652 | Orphan module 2 | Alternate toxin domain | S-type pyocin (PF06958) | 99, 9 |
| CPBP_00651 |  | Antitoxin | Colicin/pyocin immunity protein (PF01320) | 100,13 |
| ***Polymorphic Toxin System III*** |  |  |  |  |
| CPBP_00960 |  | Presumed complete multidomain toxin | *No strong match to any family or structure* |  |
| CPBP_00961 |  | Presumed antitoxin | *No strong match to any family or structure* |  |
| CPBP_00962 | Possible Orphan module 1 | Alternate toxin domain | *No strong match to any family or structure* |  |
| CPBP_00963 |  | Antitoxin | DUF3885 (PF13021) | 100, 32 |
| CPBP_00964 | Not an orphan module | - | *Short, uninformative matches against a variety of Pfam and PDB entries* |  |
| CPBP_00965 |  | - | *Short, uninformative matches against a variety of Pfam and PDB entries* |  |
| CPBP_00966 | Possible Orphan module 2 | Alternate toxin domain | Metallopeptidase toxin 5 (PF15641) | 98, 12 |
| CPBP_00967 |  | Antitoxin | Colicin-like immunity protein (PF09204) | 92,13 |
| CPBP_00968 | Possible Orphan module 3 | Alternate toxin domain | *No strong match to any family or structure* |  |
| CPBP_00969 |  | Antitoxin | *No strong match to any family or structure* |  |
| CPBP_00970 | Unlikely Orphan module 4 | Alternate toxin domain | *No strong match to any family or structure* |  |
| CPBP_00971 |  | Antitoxin | Metalloprotease (PF03410) |  |

In PTS I, the full-length toxin (CPBP_00219) terminates in an AHH nuclease domain. Accordingly, the following gene (in the coding direction) CPBP_00218 encodes a protein matching Immunity Protein 43 (Pfam: PF15570), predicted to act as antidote to the nuclease activity (67). Interestingly, it also strongly matches two Pfam families, Gmx_para_CXXCG (PF09535) and DUF1629 (PF00791), suggesting that these too may encode antitoxins. These genes are followed by two orphan modules, each containing a toxin and an antitoxin (Table 2). The next three genes encode proteins without recognisable homology to known structures or protein families, but CPBP_00210 clearly matches Immunity protein 49 (Pfam: PF15575), which is found in other PTS (67). This suggests, given their context, that CPBP_00213-CPBP_00212 and CPBP_00211-CPBP_00210 may encode third and fourth orphan modules respectively with CPBP_00211 (matching Domain of Unknown Function DUF1837; PF08878) coding for a novel toxin domain.

The full-length toxin of PTS II (encoded by CPBP_00656) contains a Tox-HNH-HHH nuclease domain (PF15637) and is followed by its cognate antitoxin encoded by CPBP_00655. Two orphan modules follow, each containing toxin-antitoxin pairs that are unambiguously homologous to well-characterised families (Table 2). However, the CPBP_00654 sequence contains no homology to the predicted N-terminal domain of the CPBP_00656 toxin; therefore, it is unclear whether the CPBP654-CPBP653 orphan module is functional.

Sequence homology in the predicted N-terminal regions of CPBP_00962, CPBP_00966, CPBP_00968, CPBP_00970 and the presumed (given its position) full-length toxin encoded by CPBP_00960 indicate the presence of a third PTS (Fig 5). Unlike PTS I and II, most predicted proteins at the PTS III locus lack recognisable domains; only CPBP_00966 and CPBP_00967 display homologies to known PTS proteins, a Metallopeptidase toxin 5 (PF15641; (67)), and Colicin-like immunity protein (PF09204) respectively. Thus, the specification of functional orphan modules is least certain for this PTS, and it may be rich in novel toxin and antitoxin genes.

RNA-seq analysis of bulk *B. saltans* cell culture confirmed that genes comprising the putative PTS systems are all transcribed (Supplementary table 2; Supplementary figure 7). A few proteins of each of these polymorphic toxin systems were also predicted as type VI secretion effector *in silico* (Supplementary table 3), suggesting the putative role of T6SS in transport and secretion of the involved toxin-antitoxin pairs.

Besides these three PTS, the endosymbiont genome also encodes for a toxin-antitoxin pair associated with T6SS. It has an anti-toxin gene (YwqK family protein), upstream of one of the VgrG genes, as previously observed (58, 68).

### Attempts to cure the symbiont result in host death

*B. saltans* cells were treated with antibiotics to cure them of endosymbiotic bacteria. Treatment with rifampicin (20 μg/ml) led to a decrease in cell count after 24 hours of rifampicin treatment compared to the untreated cells. Cell counts from three independent experiments were compared and the difference in cell growth rate was found to be statistically significant (Paired t-test: t = 5.906, d.f. = 2, p = 0.0275). By contrast, when the same treatment was applied to cell cultures of two other kinetoplastids that lack endosymbionts *(Trypanosoma theileri* and *Leptomonas costaricensis)*, rifampicin had no effect on cell number (Fig. 6).

**Figure 6:**
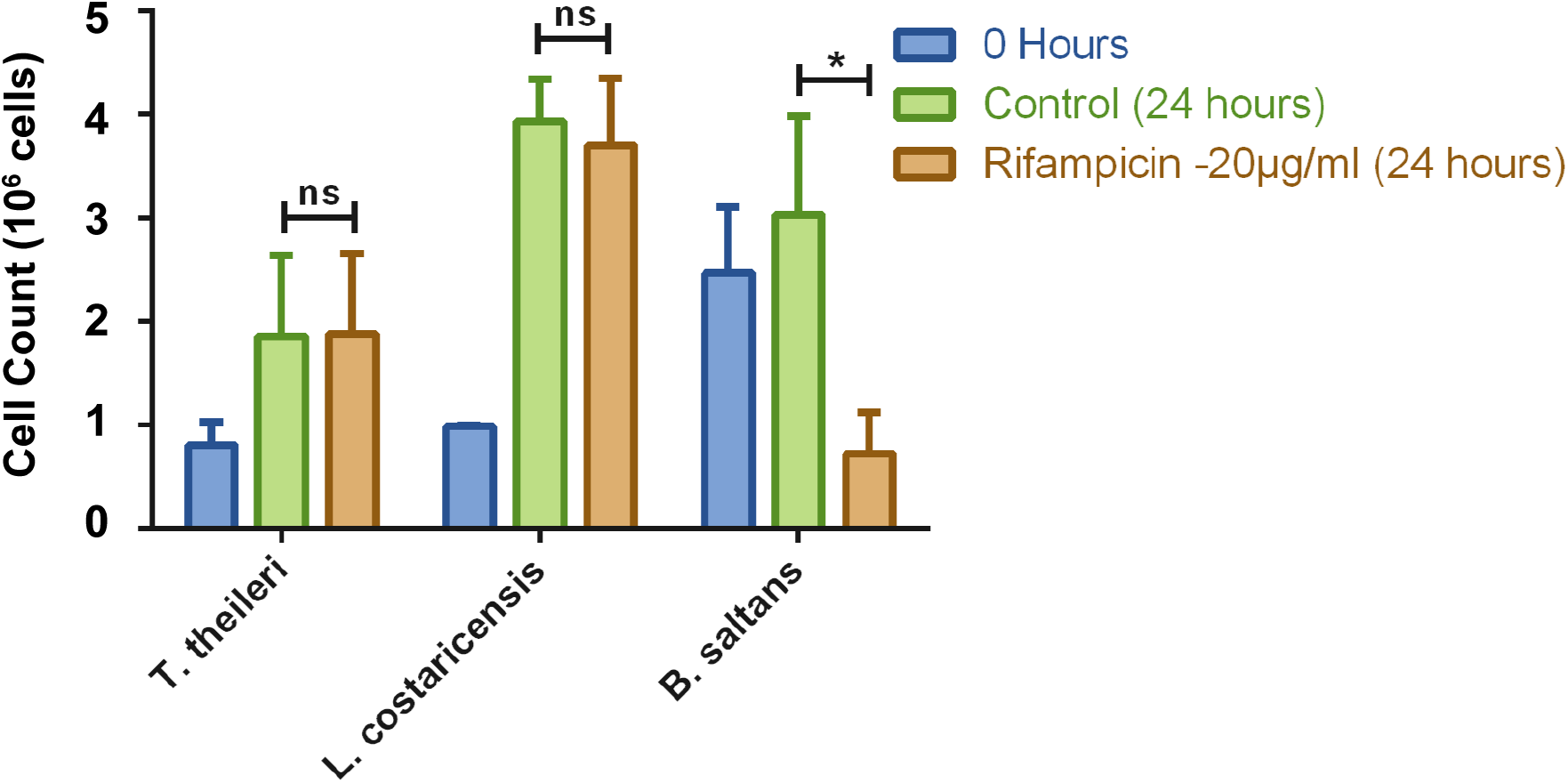
Rifampicin treatment of kinetoplastids with and without endosymbionts. Three kinetoplastids were treated with rifampicin and observed for their behavior on treatment. *B. saltans* possesses an endosymbont, *T. theileri* and *L. costaricensis* are parasitic species that do not. The graph shows number of cells before treatment and after 24 hours of treatment. The bars are average of three independent experiments. Statistical significance was calculated using ratio paired T-test.

## Discussion

The *B. saltans* endosymbiont is identified here as an alpha-proteobacterium and a novel genus of the Holosporales, *Ca*. Bodocaedibacter. The complete genome indicates that the endosymbiont is incapable of synthesizing essential amino acids, vitamins and cofactors, but instead possesses an arsenal of membrane transporters for importing essential nutrients, a specialized secretory pathway and three putative PTS operons. Antibiotic treatment of *B. saltans* results in host cell death, indicating that *Ca*. B. vickermanii is essential for host viability, despite the meagre benefits it seems to offer. Certainly, the *Bodo* endosymbiont is unrelated, phylogenetically and physiologically, to the obligate bacterial endosymbionts of parasitic kinetoplastids such as *Ca*. Kinetoplastibacterium (Alcaligenaceae) (16, 69) and *Ca*. Pandoraea (Burkholderiaceae) (15). These beta-proteobacterial symbionts can synthesize various essential amino acids independently (*Ca*. Pandoraea) (18), or in a cooperative manner with the host (*Ca*. Kinetoplastibacterium) (19), and also provide cofactors, vitamins and heme to their hosts (70). Since *Ca*. B. vickermanii lacks the genes to perform such functions, we hypothesize that host dependency in its case arises from the expression of “addictive” bacterial toxin-antitoxin proteins.

This idea is immediately plausible when we consider the ecological strategies of related endosymbionts. While *Holospora* may increase host survival under adverse conditions (71, 72), it restrains host growth in normal conditions (73, 74). An *Acanthamoeba* endosymbiont, *Ca*. Amoebophilus asiaticus, is parasitic; its genome lacks essential metabolic pathways but instead contains diverse genes shown to modulate host gene expression (59). Another *Acanthamoeba* endosymbiont, *Ca*. Jidaibacter acanthamoeba, also encodes many proteins with eukaryotic-like domains thought to interact with the host (75). Although the *Ca*. B. vickermanii genome encodes few eukaryotic-like domains, there are 339 uncharacterized proteins, most of which are lineage-specific and predicted to be secreted; such proteins could include factors for manipulating host physiology.

Perhaps the most pertinent comparison, however, is *Caedibacter*, an endosymbiont of *Paramecium* that is known for its ‘Killer trait’, which ensures its transmission at cell division (76). *Caedibacter* provides a growth advantage to *Paramecium* cells (72) but also has an adaptation to ensure its spread through the population. A portion of the endosymbiont population forms ‘R-bodies’ that are secreted outside the host cell, where they are lethal to *Paramecium* lacking the endosymbiont. In effect, the R-bodies become a toxin, against which *Caedibacter* provides protection, meaning that this is not a simple mutualism. The phylogenetic position of *Ca*. B. vickermanii among these various endosymbionts, with their ambiguous attitudes towards their hosts, makes it plausible that *Ca*. B. vickermanii too has an antagonistic mechanism for maintaining endosymbiosis.

Mechanisms like these are often described as evolved dependencies, exemplified by selfish genetic elements such as plasmids, which express a toxin-antitoxin (TA) system causing posts-egregation killing or addiction (77). TA systems in bacteria can also lead to programmed cell death (78) or persistence in response to stress conditions (79). Another evolved dependency involves the wasp *Asobara tabida* and its symbiont *Wolbachia*, which the wasp requires for oogenesis and formation of a viable offspring. As *Wolbachia* is primarily transmitted through females, loss of the same strain of *Wolbachia* in males can result in cytoplasmic incompatibility, leading to offspring mortality (80, 81). Similarly, in *C. elegans*, the peel-zeel system causes offspring to die if they do not carry the same allele as the sperm parent (12). These instances of evolved dependency on a particular allele (peel-zeel), a genetic element (plasmids), or an endosymbiont (bacteria) carrying the relevant genes are fascinating examples of addiction, where losing the addictive element can lead to host death. The presence in *Ca*. B. vickermanii of three, actively transcribed PTS and a T6SS that could convey these effector proteins, coupled with an inability to survive in an aposymbiotic state, suggest that *B. saltans* is also subject to an evolved dependency.

Symbiosis is a well-known and intricate phenomenon found at various levels of biological organization and prokaryotic endosymbionts have been instrumental in the adaptive radiation of eukaryotes (2, 82). The Family Paracaedibacteraceae is associated with diverse protists, including Rhizaria, Excavates and Amoebozoans (83–85), and now kinetoplastid flagellates. Given the diversity of their hosts, and the evolutionary distances between them, this alpha-proteobacteria lineage would seem to have been associated with eukaryotes throughout their evolutionary diversification. Like other members of the lineage, the metabolic insufficiency, specialized secretory pathway and toxin-antitoxin systems of the *Ca*. B. vickermanii genome indicate that this long relationship has not been exactly harmonious, the genomes bears witness to a struggle to retain the cooperation of their eukaryotic hosts.

## Acknowledgements

This research was supported by a Leverhulme Trust Research Grant to APJ (RPG-2014-005). We thank Alison Beckett, Department of Cellular and Molecular Physiology, for helping in electron microscopy studies. We acknowledge Liverpool Centre for Cell Imaging (CCI) for providing access to microscopy equipment and technical assistance.

