## Supplementary Figures 1-7 for "The *Paracaedibacter*-like endosymbiont of *Bodo saltans* (Kinetoplastida) uses multiple putative toxin-antitoxin systems to maintain its host association"

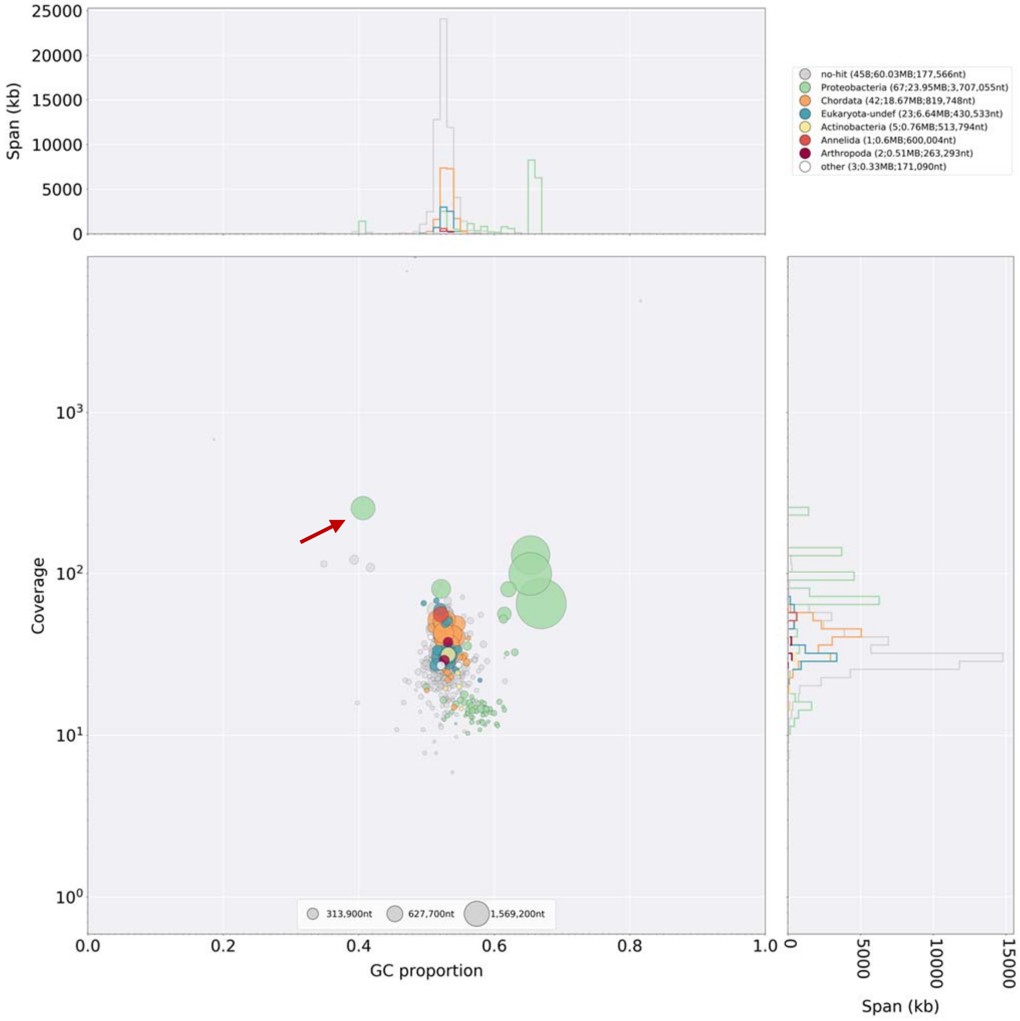


**Supplementary Figure 1: Blobplot analysis of genome assembly**

Analysis of genome assembly by mapping sequencing reads back to assembly and blast analysis against nt database of NCBI. Various contigs are plotted in the graph, with GC content on X-axis and coverage on Y-axis. Size of the circle is proportional to size of the contig. Different biological lineages are represented in different colours. Circle marked with an arrow represents the endosymbiont genome.

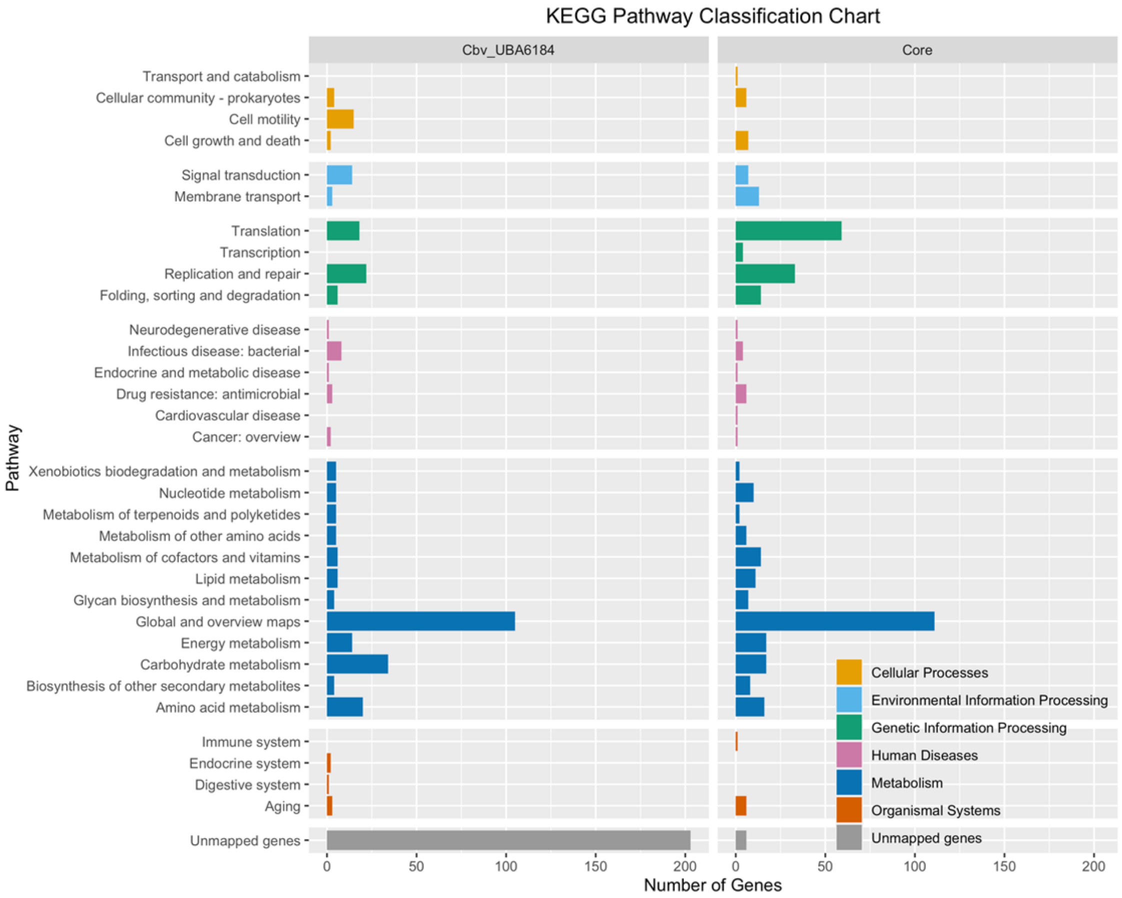


**Supplementary Figure 2: KEGG Pathway Classification Chart**

Based on the OrthoMCL analysis, core orthologues (shared by 13 protistan endosymbionts) and orthologues limited to *Candidatu*s Bodocaedibacter vickermanii (Cbv) and *Candidatus* Paracaedibacteraceae bacterium UBA6184 were identified. The bar graph shows the number of genes mapped on each pathway type on KEGG database. The colour of bars represents the six major pathway categories. Genes that did not map to any of the pathways are labelled as unmapped genes.

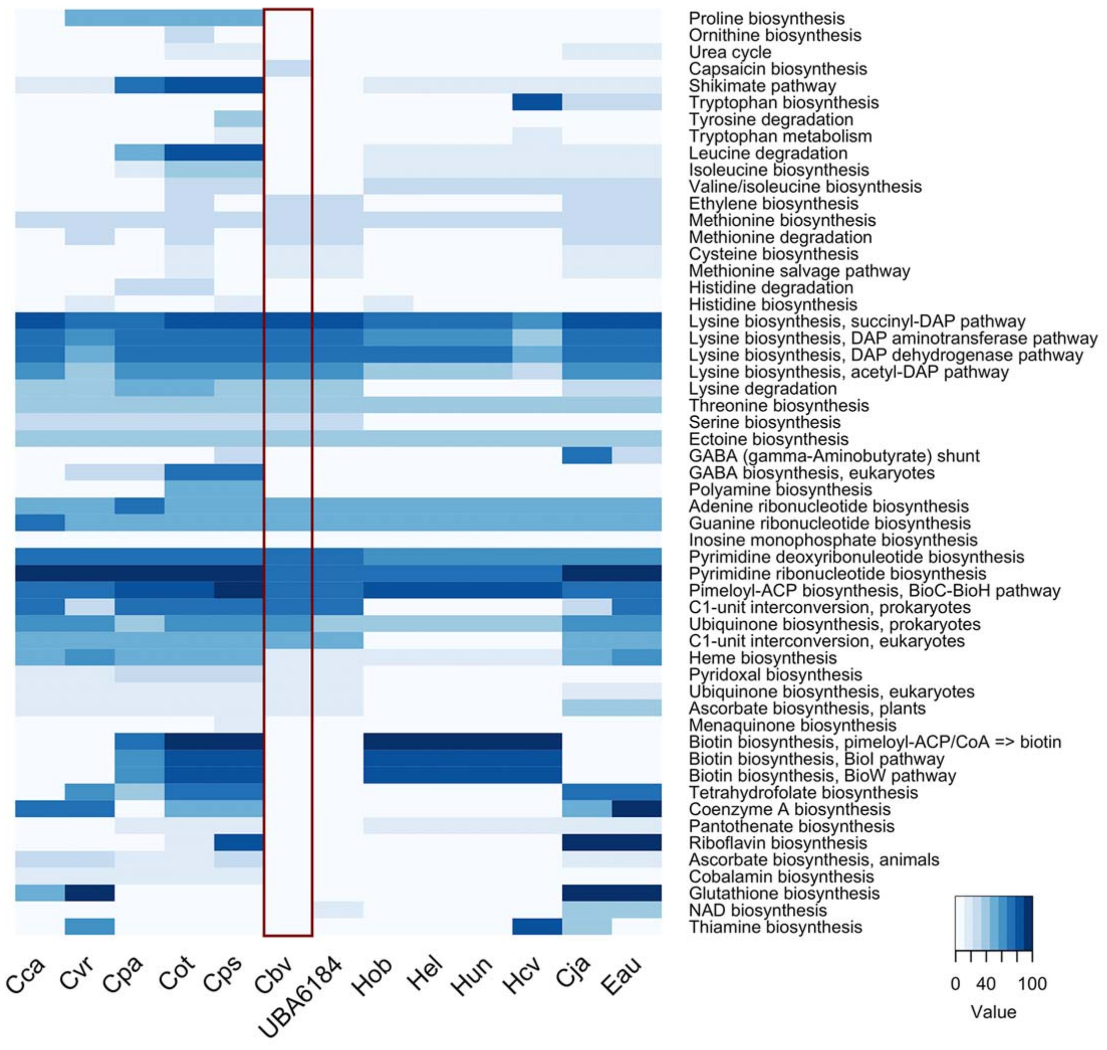


**Supplementary Figure 3: Comparison of metabolic pathways based on genome annotations for *Candidatus* Bodocaedibacter vickermanii and other alpha-proteobacterial endosymbionts of protists.**

Pathways involved in purine metabolism, pyrimidine metabolism, amino acid metabolism, polyamine biosynthesis, cofactor and vitamin biosynthesis are compared. The heatmap is based on MCR (Module completion ratio) values calculated using MAPLE v2.3.0. MCR is an indicator of completeness of a pathway in an organism. Based on the number of components of a pathway encoded by a genome, score varies from 0-100. MCR score of 100 means a complete metabolic pathway is encoded by the genome.

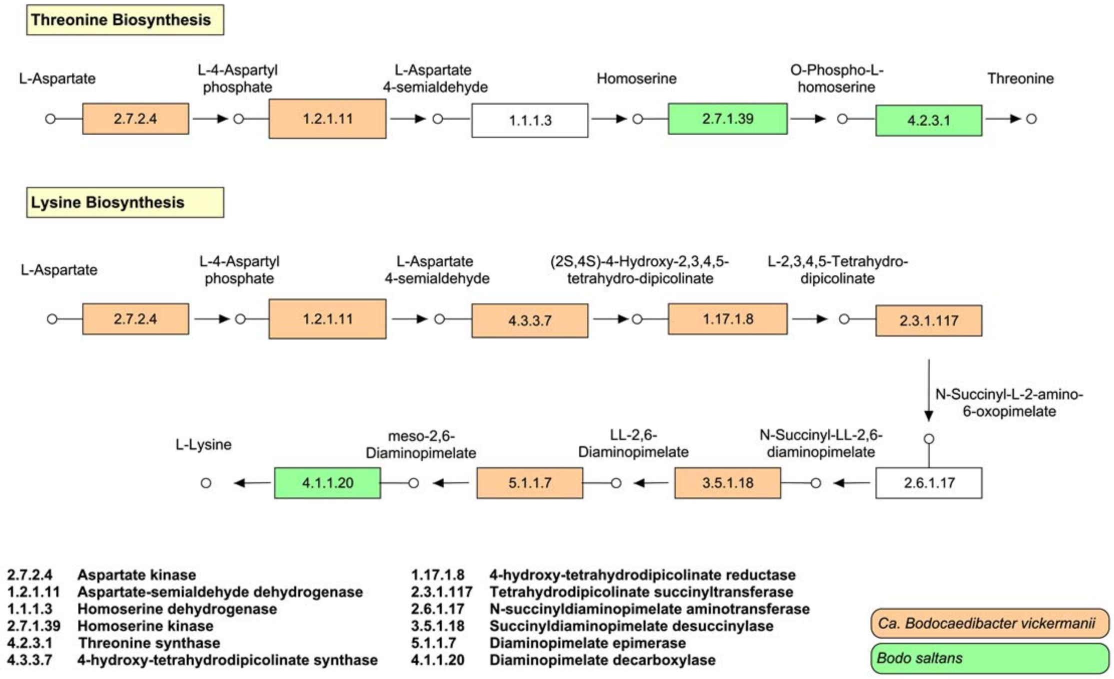


**Supplementary Figure 4: Amino acids biosynthesis pathways in *Candidatus* Bodocaedibacter vickermanii and *Bodo saltans***

Biosynthesis pathways for synthesis of L-lysine and threonine from L-aspartate are shown. Coloured boxes represent the presence of genes encoding for specific enzymes in *Ca.* B. vickermanii (in green) and *B. saltans* (in orange). White boxes represent the enzymes for which encoding genes could not be found in both organisms.

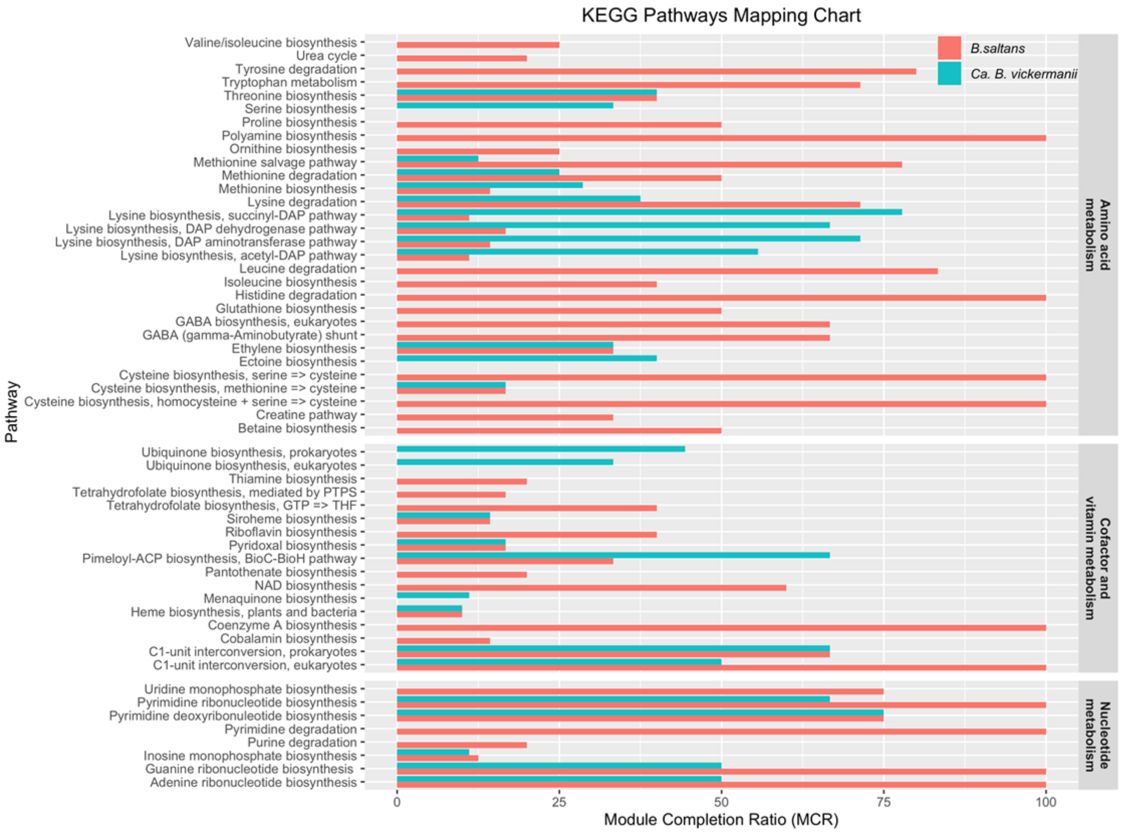


**Supplementary Figure 5: KEGG Pathway Classification Chart**

Pathways belonging to nucleotide metabolism, amino acid metabolism and cofactor and vitamins metabolism are compared between *Bodo saltans* and its endosymbiont, *Candidatu*s Bodocaedibacter vickermanii. The bar graph is based on MCR (Module completion ratio) values calculated using Genomaple v2.3.2. MCR is an indicator of completeness of a pathway in an organism. Based on the number of components of a pathway encoded by a genome, score varies from 0-100. MCR score of 100 means a complete metabolic pathway is encoded by the genome.

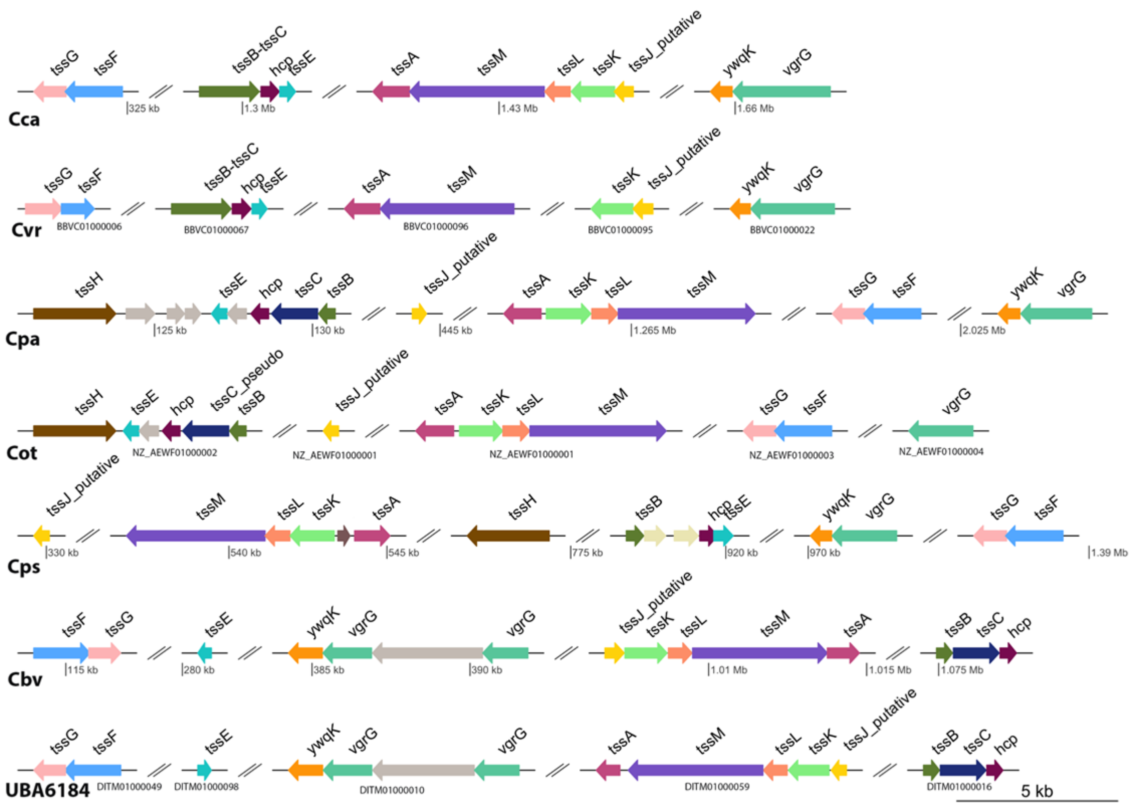


**Supplementary Figure 6: Schematic structure of Type VI Secretion System cluster in *Candidatus* Bodocaedibacter vickermanii and other alpha-proteobacterial endosymbionts of Paracedibacteraceae and Caedimonadaceae.**

T6SS core genes are dispersed at four or more locations in the genome. All maps are on the scale. Legends in grey show the location in chromosome (complete genomes) or the contig accession number (draft genomes).

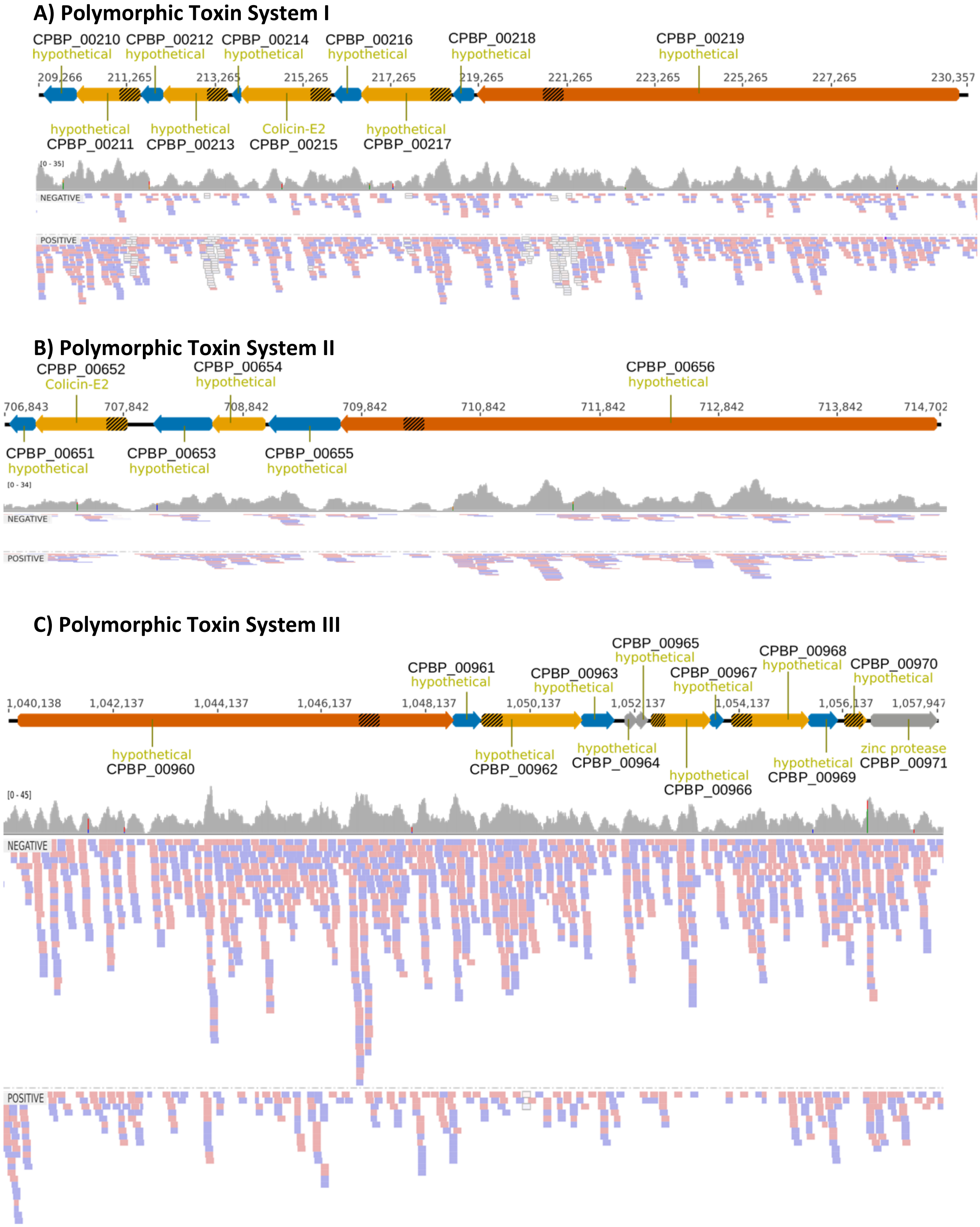


**Supplementary Figure 7: Polymorphic toxin systems in *Ca.* B. vickermanii and gene expression**

Schematic diagrams of the organization of the three predicted polymorphic toxin systems in *Ca*. B. vickermanii. RNA-seq reads are mapped on various genes and strand-specific reads quantified by SeqMonk program are shown separately.
