## Supplementary Table 1 for "The *Paracaedibacter*-like endosymbiont of *Bodo saltans* (Kinetoplastida) uses multiple putative toxin-antitoxin systems to maintain its host association"

### Supplementary Table 1: List of genes in *Candidatus* Bodocaedibacter vickermanii encoding for transporters, BLAST hits and TCDB classification

| Gene_ID | Length (amino acids) | Best Blast Hit (genBank Accession Number) | E-value | Transport Classification Database (TCDB) Family | | Number of transmenbrane α-helices |
|---|---|---|---|---|---|---|
| | | | | Family ID | Family name | |
| CPBP_00565 | 516 | ATP/ADP translocase (Fragment) Lawsonia intracellularis (B0RZB7) | 1.0E-172 | 2.A.12. | The ATP:ADP Antiporter (AAA) Family | 10 |
| CPBP_00255 | 389 | Proton/Sodium-glutamate symport protein Bacillus stearothermophilus (P24943) | 1.3E-11 | 2.A.23. | The Dicarboxylate/Amino Acid:Cation (Na+ or H+) Symporter (DAACS) Family | 9 |
| CPBP_00720 | 400 | Excitatory amino acid transporter 1 (Sodium dependent glutamate/aspartate transporter 1) Rattus norvegicus (P24942) | 7.4E-12 | 2.A.23. | The Dicarboxylate/Amino Acid:Cation (Na+ or H+) Symporter (DAACS) Family | 10 |
| CPBP_00377 | 381 | Branched-chain amino acid transport system carrier protein brnQ Chlamydia trachomatis (O84558) | 1.5E-62 | 2.A.26. | The Branched Chain Amino Acid:Cation Symporter (LIVCS) Family | 11 |
| CPBP_00197 | 434 | Arginine/agmatine antiporter Escherichia coli (P60061) | 3.8E-74 | 2.A.3. | The Amino Acid-Polyamine-Organocation (APC) Family | 11 |
| CPBP_00879 | 402 | Tyrosine-specific transport protein Escherichia coli (P0AAD4) | 1.7E-101 | 2.A.42. | The Hydroxy/Aromatic Amino Acid Permease (HAAAP) Family | 11 |
| CPBP_00324 | 300 | Uncharacterized transporter HP_1234 Helicobacter pylori (O25832) | 5.2E-25 | 2.A.7 | The Drug/Metabolite Transporter (DMT) Superfamily | 10 |
| CPBP_01233 | 271 | Hypothetical protein RP076 Rickettsia prowazekii (Q9ZE70) | 1.4E-17 | 2.A.7. | The Drug/Metabolite Transporter (DMT) Superfamily | 9 |

| | | | | | | |
|---|---|---|---|---|---|---|
| CPBP_00271 | 456 | Sodium/Pantothenate symporter (Pantothenate permease) Escherichia coli (P16256) | 4.8E-23 | 2.A.21. | The Solute:Sodium Symporter (SSS) Family | 13 |
| CPBP_01198 | 501 | Phenylacetic acid permease Pseudomonas putida (O50471) | 3.6E-163 | 2.A.21. | The Solute:Sodium Symporter (SSS) Family | 13 |
| CPBP_00044 | 227 | Major facilitator family transporter Legionella pneumophila (Q5ZUB4) | 1.1E-38 | 2.A.1. | The Major Facilitator Superfamily (MFS) | 6 |
| CPBP_00045 | 189 | Major facilitator family transporter Legionella pneumophila (Q5ZUB4) | 4.1E-37 | 2.A.1. | The Major Facilitator Superfamily (MFS) | 5 |
| CPBP_00198 | 404 | Bicyclomycin resistance protein (Sulfonamide resistance protein) Escherichia coli (P28246) | 1.4E-39 | 2.A.1. | The Major Facilitator Superfamily (MFS) | 12 |
| CPBP_00206 | 444 | Putative haloacid permease Burkholderia cepacia (Q7X4L6) | 4.8E-30 | 2.A.1. | The Major Facilitator Superfamily (MFS) | 10 |
| CPBP_00232 | 432 | Major facilitator superfamily MFS_1 Nostoc punctiforme (B2JBG5) | 2.6E-64 | 2.A.1. | The Major Facilitator Superfamily (MFS) | 10 |
| CPBP_00250 | 400 | Bicyclomycin resistance protein Escherichia coli (P28246) | 7.4E-41 | 2.A.1. | The Major Facilitator Superfamily (MFS) | 12 |
| CPBP_00330 | 419 | Proline/betaine transporter Escherichia coli (P0C0L7) | 5.3E-78 | 2.A.1. | The Major Facilitator Superfamily (MFS) | 12 |
| CPBP_00368 | 1024 | 2-acylglycerophosphoethanolamine acyltransferase Bradyrhizobium japonicum (Q89SS6) | 1.4E-57 | 2.A.1. | The Major Facilitator Superfamily (MFS) | 13 |
| CPBP_00437 | 404 | Bicyclomycin resistance protein Escherichia coli (P28246) | 2.1E-31 | 2.A.1. | The Major Facilitator Superfamily (MFS) | 12 |
| CPBP_00467 | 436 | Putative Major facilitator family transporter Legionella pneumophila (I7I571) | 4.9E-70 | 2.A.1. | The Major Facilitator Superfamily (MFS) | 12 |
| CPBP_00469 | 399 | Protein ampG Escherichia coli (P0AE16) | 1.6E-45 | 2.A.1. | The Major Facilitator Superfamily (MFS) | 12 |

| | | | | | | |
|---|---|---|---|---|---|---|
| CPBP_00470 | 414 | Putative signal transducer protein<br>Neisseria gonorrhoeae (Q5F6G0) | 6.6E-52 | 2.A.1. | The Major Facilitator Superfamily (MFS) | 10 |
| CPBP_00549 | 417 | Putative Major facilitator family transporter<br>Legionella pneumophila (I7I0I4) | 9.4E-52 | 2.A.1. | The Major Facilitator Superfamily (MFS) | 12 |
| CPBP_00557 | 264 | Putative to nasA protein<br>Legionella pneumophila (I7I1R6) | 1.4E-55 | 2.A.1. | The Major Facilitator Superfamily (MFS) | 6 |
| CPBP_00558 | 163 | Putative to nasA protein<br>Legionella pneumophila (I7I1R6) | 6.9E-16 | 2.A.1. | The Major Facilitator Superfamily (MFS) | 4 |
| CPBP_00559 | 397 | Bicyclomycin resistance protein<br>Escherichia coli (P28246) | 3.2E-36 | 2.A.1. | The Major Facilitator Superfamily (MFS) | 12 |
| CPBP_00586 | 422 | Major facilitator superfamily MFS_1<br>Nostoc punctiforme (B2JBG5) | 2.6E-71 | 2.A.1. | The Major Facilitator Superfamily (MFS) | 10 |
| CPBP_00795 | 464 | Glycerol-3-phosphate transpoter (GlpT)<br>Rickettsia prowazekii (Q9ZE92) | 9.9E-97 | 2.A.1. | The Major Facilitator Superfamily (MFS) | 12 |
| CPBP_00947 | 413 | Glycerol-3-phosphate transpoter (GlpT)<br>Rickettsia prowazekii (Q9ZE92) | 1.1E-106 | 2.A.1. | The Major Facilitator Superfamily (MFS) | 10 |
| CPBP_01035 | 416 | Putative Major facilitator family transporter<br>Legionella pneumophila (I7HQW2) | 8.9E-60 | 2.A.1. | The Major Facilitator Superfamily (MFS) | 12 |
| CPBP_01108 | 458 | Proline/betaine transporter<br>Escherichia coli (P0C0L7) | 6.5E-30 | 2.A.1. | The Major Facilitator Superfamily (MFS) | 10 |
| CPBP_01110 | 421 | Inner membrane metabolite transport protein<br>yhjE<br>Escherichia coli (P37643) | 2.1E-63 | 2.A.1. | The Major Facilitator Superfamily (MFS) | 12 |
| CPBP_00267 | 276 | Regulator of acetyl-CoA synthetase activity<br>Saccharomyces cerevisiae (P33303) | 2.0E-13 | 2.A.29. | The Mitochondrial Carrier (MC) Family | 2 |
| CPBP_00034 | 284 | ABC transporter permease protein<br>Streptococcus pyogenes (Q99ZY4) | 1.4E-38 | 3.A.1. | The ATP-binding Cassette (ABC) Superfamily | 7 |
| CPBP_00035 | 249 | Putative ABC transporter (ATP-<br>binding protein)<br>Streptococcus pyogenes (Q99ZY3) | 1.0E-51 | 3.A.1. | The ATP-binding Cassette (ABC) Superfamily | 0 |

| | | | | | | |
|---|---|---|---|---|---|---|
| CPBP_00036 | 317 | BC transporter substrate-binding protein Streptococcus pyogenes (Q99ZY6) | 5.5E-23 | 3.A.1. | The ATP-binding Cassette (ABC) Superfamily | 0 |
| CPBP_00040 | 945 | ABC transporter related Chloroflexus aurantiacus (A9WBR9) | 4.4E-11 | 3.A.1. | The ATP-binding Cassette (ABC) Superfamily | 0 |
| CPBP_00058 | 251 | Zinc import ATP-binding protein znuC Escherichia coli (P0A9X1) | 3.1E-58 | 3.A.1. | The ATP-binding Cassette (ABC) Superfamily | 0 |
| CPBP_00059 | 264 | High-affinity zinc uptake system membrane protein znuB Escherichia coli (P39832) | 2.1E-26 | 3.A.1. | The ATP-binding Cassette (ABC) Superfamily | 7 |
| CPBP_00106 | 594 | ABC transporter-related protein Ralstonia metallidurans (Q1LRE9) | 0.0E+00 | 3.A.1. | The ATP-binding Cassette (ABC) Superfamily | 6 |
| CPBP_00140 | 525 | OptrA Enterococcus faecalis (ANC59923) | 1.5E-99 | 3.A.1. | The ATP-binding Cassette (ABC) Superfamily | 0 |
| CPBP_00180 | 235 | Lipoprotein releasing system ATP-binding protein lolD Escherichia coli (P75957) | 2.8E-59 | 3.A.1. | The ATP-binding Cassette (ABC) Superfamily | 0 |
| CPBP_00181 | 416 | Lipoprotein releasing system transmembrane protein lolE Escherichia coli (P75958) | 3.7E-50 | 3.A.1. | The ATP-binding Cassette (ABC) Superfamily | 4 |
| CPBP_00283 | 614 | ABC transporter Acetobacter aceti (Q2PGB8) | 6.8E-134 | 3.A.1. | The ATP-binding Cassette (ABC) Superfamily | 0 |
| CPBP_00298 | 338 | ABC transporter substrate-binding protein Streptococcus pyogenes (Q99ZY6) | 8.7E-40 | 3.A.1. | The ATP-binding Cassette (ABC) Superfamily | 1 |
| CPBP_00307 | 560 | Energy-dependent translational throttle protein EttA Mycobacterium tuberculosis (P9WQK3) | 0.0E+00 | 3.A.1. | The ATP-binding Cassette (ABC) Superfamily | 0 |
| CPBP_00477 | 260 | Predicted ATPase involved in cell division Caldanaerobacter subterraneus (Q8R8L8) | 2.1E-47 | 3.A.1. | The ATP-binding Cassette (ABC) Superfamily | 0 |
| CPBP_00583 | 150 | Probable phospholipid ABC transporter-binding protein mlaD Escherichia coli (P64604) | 3.3E-15 | 3.A.1. | The ATP-binding Cassette (ABC) Superfamily | 1 |

| | | | | | | |
|---|---|---|---|---|---|---|
| CPBP_00784 | 371 | Putrescine-binding periplasmic protein precursor<br>Escherichia coli (P31133) | 1.4E-49 | 3.A.1. | The ATP-binding Cassette (ABC) Superfamily | 1 |
| CPBP_00785 | 375 | Putrescine transport ATP-binding protein potG<br>Escherichia coli (P31134) | 3.0E-125 | 3.A.1. | The ATP-binding Cassette (ABC) Superfamily | 0 |
| CPBP_00786 | 311 | Putrescine transport system permease protein potH<br>Escherichia coli (P31135) | 2.0E-103 | 3.A.1. | The ATP-binding Cassette (ABC) Superfamily | 6 |
| CPBP_00787 | 269 | Putrescine transport system permease protein potI<br>Escherichia coli (P0AFL1) | 5.5E-82 | 3.A.1. | The ATP-binding Cassette (ABC) Superfamily | 6 |
| CPBP_00805 | 240 | Probable amino-acid ABC transporter ATP-binding protein yqiZ<br>Bacillus subtilis (P54537) | 5.0E-97 | 3.A.1. | The ATP-binding Cassette (ABC) Superfamily | 0 |
| CPBP_00806 | 218 | Probable amino-acid ABC transporter permease protein yqiY<br>Bacillus subtilis (P54536) | 8.2E-65 | 3.A.1. | The ATP-binding Cassette (ABC) Superfamily | 3 |
| CPBP_00807 | 256 | Probable amino-acid ABC transporter extracellular-binding protein yqiX<br>Bacillus subtilis (P54535) | 2.6E-36 | 3.A.1. | The ATP-binding Cassette (ABC) Superfamily | 0 |
| CPBP_00918 | 253 | Hypothetical protein At1g19800 (Permease-like protein)<br>Arabidopsis thaliana (Q8L4R0) | 2.3E-46 | 3.A.1. | The ATP-binding Cassette (ABC) Superfamily | 5 |
| CPBP_00919 | 252 | Putative ATPase component of ABC transporter system<br>Sphingobium japonicum (A4PCH8) | 2.1E-53 | 3.A.1. | The ATP-binding Cassette (ABC) Superfamily | 0 |
| CPBP_01091 | 269 | High-affinity zinc uptake system protein znuA precursor<br>Escherichia coli (P39172) | 5.6E-21 | 3.A.1. | The ATP-binding Cassette (ABC) Superfamily | 0 |
| CPBP_01190 | 596 | Mitochondrial ATP-binding cassette 2<br>Homo sapiens (Q9NRK6) | 1.8E-108 | 3.A.1. | The ATP-binding Cassette (ABC) Superfamily | 5 |
| CPBP_01236 | 519 | Mitochondrial transpoter ATM1<br>Rickettsia prowazekii (Q9ZDW0) | 3.2E-30 | 3.A.1. | The ATP-binding Cassette (ABC) Superfamily | 5 |
| CPBP_00691 | 315 | Putative uncharacterized protein<br>Hoeflea phototrophica (A9DHE6) | 2.4E-52 | 2.A.83. | The Na+-dependent Bicarbonate Transporter (SBT) Family | 8 |

| | | | | | | |
|---|---|---|---|---|---|---|
| CPBP_00050 | 178 | BioY family protein<br>Rickettsia typhi (Q68X47) | 7.7E-20 | 2.A.88. | The Vitamin Uptake Transporter (VUT or ECF) Family | 5 |
| CPBP_00587 | 1030 | Probable Resistance-Nodulation- Cell Division (RND) efflux transporter<br>Pseudomonas aeruginosa (Q9HW27) | 0.0E+00 | 2.A.6. | The Resistance-Nodulation-Cell Division (RND) Superfamily | 12 |
| CPBP_00588 | 382 | Probable RND efflux membrane- fusion protein<br>Burkholderia glumae (Q4VSJ3) | 2.9E-41 | 2.A.6. | The Resistance-Nodulation-Cell Division (RND) Superfamily | 1 |
| CPBP_00823 | 534 | Protein-export membrane protein secD<br>Escherichia coli (P0AG90) | 6.1E-86 | 2.A.6. | The Resistance-Nodulation-Cell Division (RND) Superfamily | 5 |
| CPBP_00824 | 310 | Protein translocase subunit SecF<br>Escherichia coli (P0AG93) | 5.6E-55 | 2.A.6. | The Resistance-Nodulation-Cell Division (RND) Superfamily | 6 |
| CPBP_00861 | 367 | Multidrug efflux transporter VexE<br>Vibrio cholerae (A6P7H2) | 6.5E-47 | 2.A.6. | The Resistance-Nodulation-Cell Division (RND) Superfamily | 1 |
| CPBP_00863 | 1055 | Multidrug efflux transporter VexF<br>Vibrio cholerae (A6P7H3) | 0.0E+00 | 2.A.6. | The Resistance-Nodulation-Cell Division (RND) Superfamily | 12 |
| CPBP_00312 | 514 | Virulence factor mviN<br>Salmonella typhimurium (P37169) | 1.2E-66 | 2.A.66. | The Multidrug/Oligosaccharidyl-lipid/Polysaccharide (MOP) Flippase Superfamily | 13 |
| CPBP_00313 | 524 | Virulence factor mviN<br>Salmonella typhimurium (P37169) | 3.6E-73 | 2.A.66. | The Multidrug/Oligosaccharidyl-lipid/Polysaccharide (MOP) Flippase Superfamily | 12 |
| CPBP_00898 | 455 | Probable multidrug resistance protein NorM<br>Thermotoga maritima (Q9WZS2) | 2.3E-14 | 2.A.66. | The Multidrug/Oligosaccharidyl-lipid/Polysaccharide (MOP) Flippase Superfamily | 12 |
| CPBP_00899 | 452 | Probable multidrug resistance protein NorM<br>Thermotoga maritima (Q9WZS2) | 2.4E-19 | 2.A.66. | The Multidrug/Oligosaccharidyl-lipid/Polysaccharide (MOP) Flippase Superfamily | 12 |
| CPBP_00207 | 315 | Na+/H+ antiporter<br>Vibrio parahaemolyticus (Q56725) | 1.6E-67 | 2.A.33. | The NhaA Na+:H+ Antiporter (NhaA) Family | 9 |
| CPBP_00571 | 441 | Na+/H+ antiporter family protein<br>Shewanella oneidensis (Q8EHX2) | 1.7E-127 | 2.A.35. | The NhaC Na+:H+ Antiporter (NhaC) Family | 9 |

| | | | | | | |
|---|---|---|---|---|---|---|
| CPBP_00707 | 402 | K(+) efflux antiporter 2, chloroplastic Arabidopsis thaliana (O65272) | 1.1E-91 | 2.A.37. | The Monovalent Cation:Proton Antiporter-2 (CPA2) Family | 11 |
| CPBP_00342 | 458 | Trk system potassium uptake protein trkA Escherichia coli (P0AGI8) | 2.4E-69 | 2.A.38 | The K+ Transporter (Trk) Family | 0 |
| CPBP_00990 | 483 | Trk system potassium uptake protein TrkI Halomonas elongata (Q6T3V6) | 9.3E-112 | 2.A.38. | The K+ Transporter (Trk) Family | 12 |
| CPBP_00433 | 694 | H+ translocating pyrophosphate synthase Rhodospirillum rubrum (O68460) | 0.0E+00 | 3.A.10. | The H+-translocating Pyrophosphatase (H+-PPase) Family | 13 |
| CPBP_00608 | 492 | NADH dehydrogenase 1, chain 14 Paracoccus denitrificans (P29926) | 6.5E-74 | 3.D.1. | The Proton-translocating NADH Dehydrogenase (NDH) Family | 12 |
| CPBP_00609 | 490 | NADH dehydrogenase 1, chain 13 Paracoccus denitrificans (P29925) | 7.2E-135 | 3.D.1. | The Proton-translocating NADH Dehydrogenase (NDH) Family | 14 |
| CPBP_00610 | 641 | NADH dehydrogenase 1, chain 12 Paracoccus denitrificans (P29924) | 9.4E-155 | 3.D.1. | The Proton-translocating NADH Dehydrogenase (NDH) Family | 16 |
| CPBP_00614 | 335 | NADH dehydrogenase 1, chain 8 Paracoccus denitrificans (P29920) | 3.2E-105 | 3.D.1. | The Proton-translocating NADH Dehydrogenase (NDH) Family | 8 |
| CPBP_00134 | 264 | Ion transport protein Arcobacter butzleri (A8EVM5) | 8.2E-33 | 1.A.1. | The Voltage-gated Ion Channel (VIC) Superfamily | 6 |
| CPBP_00358 | 426 | Mg2+ and Co2+ transporter CorB, contains DUF21, CBS pair, and CorC-HlyC domains Pseudomonas bauzanensis (SES26846) | 1.1E-67 | 1.A.112 | The Cyclin M Mg2+ Exporter (CNNM) Family | 3 |
| CPBP_00231 | 458 | Magnesium transporter MgtE Prochlorococcus marinus (A2C579) | 7.0E-50 | 1.A.26 | The Mg2+ Transporter-E (MgtE) Family | 5 |
| CPBP_01170 | 142 | Biopolymer transport protein exbD Escherichia coli (P0ABV2) | 1.0E-30 | 1.A.30 | The H+- or Na+-translocating Bacterial Flagellar Motor/ExbBD Outer Membrane Transport Energizer (Mot-Exb) Superfamily | 1 |
| CPBP_01171 | 223 | Protein tolQ Escherichia coli (P0ABU9) | 1.8E-52 | 1.A.30 | The H+- or Na+-translocating Bacterial Flagellar Motor/ExbBD Outer Membrane Transport Energizer (Mot-Exb) Superfamily | 3 |

| | | | | | | |
|---|---|---|---|---|---|---|
| CPBP_00693 | 547 | Membrane protein, UPF0126<br>Gramella forsetii (A0M015) | 3.9E-11 | 1.A.62. | The Homotrimeric Cation Channel (TRIC) Family | 7 |
| CPBP_00694 | 539 | Uncharacterized protein<br>Yersinia pestis (I7MT28) | 1.9E-12 | 1.A.62. | The Homotrimeric Cation Channel (TRIC) Family | 7 |
| CPBP_01086 | 482 | Putative outer membrane secretion protein<br>Rhizobium meliloti (Q92Q38) | 2.4E-26 | 1.B.17. | The Outer Membrane Factor (OMF) Family | 0 |
| CPBP_01082 | 276 | Hemolysin C<br>Brachyspira hyodysenteriae (Q54318) | 2.9E-36 | 1.C.126. | The HlyC (HlyC) Family of Haemolysins | 0 |
| CPBP_01232 | 295 | Ferrous-iron efflux pump fieF<br>Escherichia coli (P69380) | 6.1E-62 | 2.A.4. | The Cation Diffusion Facilitator (CDF) Family | 5 |
| CPBP_00925 | 369 | Permease<br>Brucella abortus (C9VRY8) | 1.6E-67 | 2.A.86. | The Autoinducer-2 Exporter (AI-2E) Family | 8 |
| CPBP_01020 | 561 | Inner membrane protein YidC<br>Escherichia coli (P25714) | 4.9E-60 | 2.A.9. | The Cytochrome Oxidase Biogenesis (Oxa1) Family | 5 |
| CPBP_00098 | 445 | Preprotein translocase subunit secY<br>Escherichia coli (P0AGA2) | 2.9E-140 | 3.A.5. | The General Secretory Pathway (Sec) Family | 10 |
| CPBP_00794 | 652 | 4-coumarate--CoA ligase 1<br>Arabidopsis thaliana (Q42524) | 2.9E-06 | 4.C.1. | The Proposed Fatty Acid Transporter (FAT) Family | 0 |
| CPBP_00837 | 587 | Ferrous iron transport protein B<br>Leptospira biflexa (Q5XPH7) | 6.0E-87 | 9.A.8. | The Ferrous Iron Uptake (FeoB) Family | 9 |
