## Supplementary Table 2 for "The *Paracaedibacter*-like endosymbiont of *Bodo saltans* (Kinetoplastida) uses multiple putative toxin-antitoxin systems to maintain its host association"

### Supplementary Table 2: List of genes in *Candidatus* Bodocaedibacter vickermanii and their expression analysis

| | | gene length | Read count | RPK values |
|---|---|---|---|---|
| **Type VI Secretion System** | | | | |
| **CPBP_00125** | Type VI secretion system baseplate subunit TssF | 1785 | 21 | 11.8 |
| **CPBP_00126** | Type VI secretion system baseplate subunit TssG | 1017 | 36 | 35.4 |
| **CPBP_00258** | type VI secretion system baseplate subunit TssE | 432 | 12 | 27.8 |
| **CPBP_00353** | type VI secretion system tip protein VgrG | 1545 | 62 | 40.1 |
| **CPBP_00355** | type VI secretion system tip protein VgrG | 1434 | 43 | 30.0 |
| **CPBP_00565** | ADP,ATP carrier protein 1 | 1551 | 214 | 138.0 |
| **CPBP_00933** | type VI secretion system protein, membrane lipoprotein | 615 | 18 | 29.3 |
| **CPBP_00934** | type VI secretion system baseplate subunit TssK | 1341 | 32 | 23.9 |
| **CPBP_00935** | type VI secretion system protein TssL (DotU family) | 759 | 18 | 23.7 |
| **CPBP_00936** | type VI secretion system membrane subunit TssM | 4272 | 133 | 31.1 |
| **CPBP_00937** | type VI secretion system protein TssA | 1020 | 46 | 45.1 |
| **CPBP_00985** | type VI secretion system contractile sheath small subunit, TssB | 513 | 22 | 42.9 |
| **CPBP_00986** | type VI secretion system contractile sheath large subunit, TssC | 1455 | 21 | 14.4 |
| **CPBP_00987** | Hcp1-like superfamily protein | 525 | 29 | 55.2 |
| **CPBP_01201** | DUF4280 domain-containing protein | 393 | 0 | 0.0 |
| **Toxin-Antitoxin System 1** | | | | |
| **CPBP_00210** | hypothetical protein | 735 | 50 | 68.0 |
| **CPBP_00211** | hypothetical protein | 1455 | 131 | 90.0 |
| **CPBP_00212** | hypothetical protein | 474 | 19 | 40.1 |
| **CPBP_00213** | hypothetical protein | 1473 | 59 | 40.1 |
| **CPBP_00214** | hypothetical protein | 207 | 5 | 24.2 |
| **CPBP_00215** | Colicin-E2 | 2043 | 65 | 31.8 |
| **CPBP_00216** | hypothetical protein | 597 | 19 | 31.8 |
| **CPBP_00217** | DUF4157 domain-containing protein | 2043 | 76 | 37.2 |
| **CPBP_00218** | hypothetical protein | 486 | 14 | 28.8 |
| **CPBP_00219** | type IV secretion protein Rhs | 10938 | 384 | 35.1 |
| **Toxin-Antitoxin System 2** | | | | |
| **CPBP_00651** | hypothetical protein | 216 | 1 | 4.6 |
| **CPBP_00652** | Colicin-E2 | 771 | 9 | 11.7 |
| **CPBP_00653** | hypothetical protein | 486 | 21 | 43.2 |
| **CPBP_00654** | hypothetical protein | 450 | 34 | 75.6 |
| **CPBP_00655** | hypothetical protein | 597 | 5 | 8.4 |
| **CPBP_00656** | hypothetical protein | 5016 | 131 | 26.1 |
| **Toxin-Antitoxin System 3** | | | | |
| **CPBP_00960** | MafB2 adhesin | 8352 | 384 | 46.0 |
| **CPBP_00961** | hypothetical protein | 519 | 36 | 69.4 |
| **CPBP_00962** | hypothetical protein | 1893 | 117 | 61.8 |
| **CPBP_00963** | hypothetical protein | 618 | 10 | 16.2 |
| **CPBP_00964** | serine kinase | 228 | 26 | 114.0 |
| **CPBP_00965** | serine kinase | 204 | 8 | 39.2 |
| **CPBP_00966** | RHS repeat-associated core domain-containing protein | 1140 | 50 | 43.9 |
| **CPBP_00967** | hypothetical protein | 249 | 16 | 64.3 |
| **CPBP_00968** | hypothetical protein | 1476 | 69 | 46.7 |
| **CPBP_00969** | hypothetical protein | 534 | 25 | 46.8 |
| **CPBP_00970** | hypothetical protein | 411 | 25 | 60.8 |
| **CPBP_00971** | putative zinc protease | 1257 | 46 | 36.6 |
