## Supplementary Table 3 for "The *Paracaedibacter*-like endosymbiont of *Bodo saltans* (Kinetoplastida) uses multiple putative toxin-antitoxin systems to maintain its host association"

### Supplementary Table 3: List of genes in *Candidatus* Bodocaedibacter vickermanii encoding for putative Type VI secretion effectors

| S. No. | Gene ID | Product | T6SE | Note | Prediction by other methods |
|---|---|---|---|---|---|
| 1 | CPBP_00008 | hypothetical protein | Yes | | SignalP and SecretomeP |
| 2 | CPBP_00009 | ATP-dependent RNA helicase DbpA | Yes | | SecretomeP |
| 3 | CPBP_00012 | peptidase | Yes | | |
| 4 | CPBP_00027 | CDGSH iron-sulfur domain-containing protein | Yes | | SecretomeP |
| 5 | CPBP_00039 | mannosyltransferase 1 family protein | Yes | | |
| 6 | CPBP_00062 | hypothetical protein | Yes | | SignalP and SecretomeP |
| 7 | CPBP_00129 | Flagellar hook-length control protein FliK | Yes | | SecretomeP |
| 8 | CPBP_00131 | Flagellar hook protein FlgE | Yes | | SecretomeP |
| 9 | CPBP_00132 | Flagellar hook protein FlgE | Yes | | SignalP and SecretomeP |
| 10 | CPBP_00138 | hypothetical protein | Yes | | SecretomeP |
| 11 | CPBP_00143 | Single-stranded DNA-binding protein | Yes | | SecretomeP |
| 12 | CPBP_00151 | RNA-splicing ligase RtcB | Yes | | |
| 13 | CPBP_00156 | phage portal protein | Yes | | |
| 14 | CPBP_00163 | phage major capsid protein | Yes | | SecretomeP |
| 15 | CPBP_00166 | hypothetical protein | Yes | | SecretomeP |
| 16 | CPBP_00168 | hypothetical protein | Yes | | |
| 17 | CPBP_00169 | hypothetical protein | Yes | | |
| 18 | CPBP_00174 | hypothetical protein | Yes | | |
| 19 | CPBP_00184 | hypothetical protein | Yes | | SecretomeP |
| 20 | CPBP_00193 | DsbA family protein | Yes | | SignalP and SecretomeP |
| 21 | CPBP_00211 | hypothetical protein | Yes | Toxin-Antitoxin system-1 (210:219) | SecretomeP |
| 22 | CPBP_00212 | hypothetical protein | Yes | | |
| 23 | CPBP_00213 | hypothetical protein | Yes | | SecretomeP |
| 24 | CPBP_00215 | Colicin-E2 | Yes | | |
| 25 | CPBP_00217 | DUF4157 domain-containing protein | Yes | | |
| 26 | CPBP_00220 | hypothetical protein | Yes | | SecretomeP |
| 27 | CPBP_00222 | Ribonuclease YobL | Yes | | SignalP |
| 28 | CPBP_00244 | Cellulosome-anchoring protein | Yes | | SecretomeP |
| 29 | CPBP_00247 | hypothetical protein | Yes | | SignalP and SecretomeP |
| 30 | CPBP_00257 | hypothetical protein | Yes | T6SS (258) | SignalP and SecretomeP |
| 31 | CPBP_00270 | TIGR02452 family protein | Yes | | SignalP and SecretomeP |
| 32 | CPBP_00289 | hypothetical protein | Yes | | SignalP |
| 33 | CPBP_00290 | hypothetical protein | Yes | | SecretomeP |

| | | | | | |
|---|---|---|---|---|---|
| 34 | CPBP_00297 | hypothetical protein | Yes | | SecretomeP |
| 35 | CPBP_00315 | hypothetical protein | Yes | | SecretomeP |
| 36 | CPBP_00326 | filamentous hemagglutinin N-terminal domain-containing protein | Yes | | |
| 37 | CPBP_00329 | Murein hydrolase activator NlpD | Yes | | SecretomeP |
| 38 | CPBP_00332 | hypothetical protein | Yes | | SignalP and SecretomeP |
| 39 | CPBP_00347 | prepilin-type N-terminal cleavage/methylation domain-containing protein | Yes | | SecretomeP |
| 40 | CPBP_00348 | type II secretion system protein | Yes | | |
| 41 | CPBP_00352 | toxin-antitoxin system, YwqK family antitoxin | Yes | | SecretomeP |
| 42 | CPBP_00353 | type VI secretion system tip protein VgrG | Yes | T6SS | SecretomeP |
| 43 | CPBP_00354 | hypothetical protein | Yes | | SecretomeP |
| 44 | CPBP_00355 | type VI secretion system tip protein VgrG | Yes | T6SS | |
| 45 | CPBP_00357 | hypothetical protein | Yes | | SignalP and SecretomeP |
| 46 | CPBP_00363 | Aconitate hydratase A | Yes | | |
| 47 | CPBP_00382 | hypothetical protein | Yes | | SignalP and SecretomeP |
| 48 | CPBP_00396 | hypothetical protein | Yes | | SecretomeP |
| 49 | CPBP_00413 | hypothetical protein | Yes | | SignalP and SecretomeP |
| 50 | CPBP_00419 | DDE transposase | Yes | | SecretomeP |
| 51 | CPBP_00420 | Chaperone protein DnaJ | Yes | | SecretomeP |
| 52 | CPBP_00430 | DUF2282 domain-containing protein | Yes | | SignalP and SecretomeP |
| 53 | CPBP_00432 | hypothetical protein | Yes | | SecretomeP |
| 54 | CPBP_00441 | hypothetical protein | Yes | | SignalP and SecretomeP |
| 55 | CPBP_00442 | hypothetical protein | Yes | | SignalP and SecretomeP |
| 56 | CPBP_00452 | Dipeptidyl aminopeptidase BIII | Yes | | |
| 57 | CPBP_00473 | hypothetical protein | Yes | | SecretomeP |
| 58 | CPBP_00484 | hypothetical protein | Yes | | SecretomeP |
| 59 | CPBP_00528 | zinc-ribbon domain-containing protein | Yes | | |
| 60 | CPBP_00535 | hypothetical protein | Yes | | SecretomeP |
| 61 | CPBP_00573 | 50S ribosomal protein L27 | Yes | | SecretomeP |
| 62 | CPBP_00582 | NADH dehydrogenase | Yes | | SecretomeP |
| 63 | CPBP_00638 | flagellar hook protein FliD | Yes | | |
| 64 | CPBP_00646 | hypothetical protein | Yes | | SignalP and SecretomeP |
| 65 | CPBP_00652 | Colicin-E2 | Yes | Toxin-Antitoxin | SecretomeP |
| 66 | CPBP_00653 | hypothetical protein | Yes | system-2 | |
| 67 | CPBP_00654 | hypothetical protein | Yes | (651:656) | |
| 68 | CPBP_00663 | hypothetical protein | Yes | | SignalP and SecretomeP |
| 69 | CPBP_00667 | hypothetical protein | Yes | | SecretomeP |
| 70 | CPBP_00685 | hypothetical protein | Yes | | SignalP and SecretomeP |
| 71 | CPBP_00696 | AraC family transcriptional regulator | Yes | | SecretomeP |
| 72 | CPBP_00697 | Vitamin B12-dependent ribonucleoside-diphosphate reductase | Yes | | |

| | | | | | |
|---|---|---|---|---|---|
| 73 | CPBP_00761 | hypothetical protein | Yes | | |
| 74 | CPBP_00764 | Murein DD-endopeptidase MepM | Yes | | SecretomeP |
| 75 | CPBP_00767 | Tol-Pal system protein TolB | Yes | | |
| 76 | CPBP_00773 | Poly(3-hydroxyalkanoate) polymerase subunit PhaC | Yes | | SecretomeP |
| 77 | CPBP_00782 | 50S ribosomal protein L31 | Yes | | SecretomeP |
| 78 | CPBP_00792 | 2-oxoglutarate dehydrogenase E1 component | Yes | | |
| 79 | CPBP_00796 | hypothetical protein | Yes | | SecretomeP |
| 80 | CPBP_00797 | hypothetical protein | Yes | | SecretomeP |
| 81 | CPBP_00798 | hypothetical protein | Yes | | SecretomeP |
| 82 | CPBP_00799 | putative transcriptional regulatory protein | Yes | | |
| 83 | CPBP_00810 | Putative penicillin-binding protein PbpX | Yes | | SignalP and SecretomeP |
| 84 | CPBP_00842 | DUF3576 domain-containing protein | Yes | | SignalP |
| 85 | CPBP_00843 | Leucine--tRNA ligase | Yes | | SecretomeP |
| 86 | CPBP_00854 | Flagellar basal-body rod protein FlgF | Yes | | SecretomeP |
| 87 | CPBP_00880 | hypothetical protein | Yes | | SecretomeP |
| 88 | CPBP_00900 | hypothetical protein | Yes | | |
| 89 | CPBP_00932 | hypothetical protein | Yes | T6SS (933:937) | SecretomeP |
| 90 | CPBP_00943 | Superoxide dismutase [Fe] | Yes | | SecretomeP |
| 91 | CPBP_00951 | Virginiamycin A acetyltransferase | Yes | | |
| 92 | CPBP_00962 | hypothetical protein | Yes | Toxin-Antitoxin system-3 (960:971) | |
| 93 | CPBP_00975 | hypothetical protein | Yes | | SignalP and SecretomeP |
| 94 | CPBP_00987 | Hcp1-like superfamily protein | Yes | T6SS (985:987) | |
| 95 | CPBP_00991 | Co2+/Mg2+ efflux protein ApaG | Yes | | SecretomeP |
| 96 | CPBP_00998 | Cysteine--tRNA ligase | Yes | | |
| 97 | CPBP_01003 | hypothetical protein | Yes | | SecretomeP |
| 98 | CPBP_01023 | hypothetical protein | Yes | | SecretomeP |
| 99 | CPBP_01026 | hypothetical protein | Yes | | SecretomeP |
| 100 | CPBP_01049 | hypothetical protein | Yes | | SecretomeP |
| 101 | CPBP_01060 | Exodeoxyribonuclease III | Yes | | |
| 102 | CPBP_01063 | hypothetical protein | Yes | | SignalP and SecretomeP |
| 103 | CPBP_01064 | hypothetical protein | Yes | | SecretomeP |
| 104 | CPBP_01071 | Penicillin-binding protein 1F | Yes | | SecretomeP |
| 105 | CPBP_01134 | hypothetical protein | Yes | | SignalP and SecretomeP |
| 106 | CPBP_01139 | putative exported protein | Yes | | SecretomeP |
| 107 | CPBP_01149 | hypothetical protein | Yes | | SignalP |
| 108 | CPBP_01150 | hypothetical protein | Yes | | SecretomeP |

| | | | | |
|---|---|---|---|---|
| 109 | CPBP_01162 | fucose permease | Yes | |
| 110 | CPBP_01180 | hypothetical protein | Yes | SignalP and SecretomeP |
| 111 | CPBP_01186 | hypothetical protein | Yes | SecretomeP |
| 112 | CPBP_01191 | hypothetical protein | Yes | SecretomeP |
| 113 | CPBP_01193 | DDE transposase | Yes | SecretomeP |
| 114 | CPBP_01194 | hypothetical protein | Yes | SignalP and SecretomeP |
| 115 | CPBP_01235 | hypothetical protein | Yes | SecretomeP |
| 116 | CPBP_01258 | outer membrane protein assembly factor BamE | Yes | SignalP and SecretomeP |
| 117 | CPBP_01259 | hypothetical protein | Yes | |
